# Computational insights into the interaction between Topoisomerase I and Rpc82 subunit of RNA Polymerase III in *Saccharomyces cereviseae*

**DOI:** 10.64898/2026.01.31.703072

**Authors:** Prakash Nandi, Izaz Monir Kamal, Saikat Chakrabarti, Sugopa Sengupta

**Author notes:** Corresponding author, Sugopa Sengupta, Co corresponding author, Saikat Chakrabarti. **E-mail IDs of other authors:** Prakash Nandi, Izaz Monir Kamal.

## Abstract

The process of DNA transcription leads to the generation of torsional stress, which must be resolved for smooth progression of the transcription machinery. In *Saccharomyces cerevisiae*, DNA topoisomerase I (Top1), a type IB topoisomerase, plays a critical role in relaxing supercoils and mitigating the topological strain associated with transcription. While several proteins from the transcription machinery have been reported to interact with yeast Top1, detailed characterization and functional relevance of these interactions have remained underexplored. This gap is partly due to the absence of a complete three-dimensional structure of the full-length enzyme, which hinders structure-based computational analyses of its interactome. In this study, we present a template-based model of full-length yeast Top1. Leveraging this model, we investigated its molecular interaction with Rpc82, a key subunit of RNA polymerase III enzyme, responsible for transcribing small non-coding RNAs such as tRNAs and 5S rRNA. Through molecular docking and molecular dynamics simulations, critical residues at the Top1–Rpc82 interface were identified that likely mediate their interaction. Our findings provide new insights into the structural basis of Top1’s association with RNA polymerase III and its potential role in regulating Pol III-mediated transcription. The Top1 model developed here offers a valuable framework for future *in silico* studies aimed at elucidating the broader interactome and regulatory mechanisms of this essential enzyme.

## 1. INTRODUCTION

The maintenance of eukaryotic genomic DNA in a negatively supercoiled state within the nucleus presents unique topological challenges during essential cellular processes like DNA replication and transcription. These challenges are dynamically regulated by the coordinated action of enzymes called DNA topoisomerases. During transcription, RNA polymerase generates excess positive supercoils ahead of the transcription bubble and negative supercoils behind, that must be relaxed by DNA topoisomerases to ensure smooth progression of the transcription machinery along the DNA template [1], [2]. RNA polymerase often gets hindered owing to accumulation of excess DNA supercoils and formation of R-loops (DNA-RNA hybrids) as inevitable tangles [3], [4], [5]. The level of supercoiling has direct effect on promoter accessibility to transcription factors and RNA polymerase binding [6]. Topoisomerase I (Top1) is a type IB topoisomerase [7], [8] which cleaves only one DNA strand to relax both positive and negative supercoils of DNA helix, help to resolve the topological constraints during transcription [8], [9], [10], [11], [12], [13]. In addition to its intrinsic catalytic role, Top1 interacts with a diverse array of proteins that either regulate its catalytic function or facilitate its precise recruitment to the regions of topological stress within the genome. For example, Top1 in budding yeast physically interacts with Csm3-Tof1 complex to regulate replication fork pausing [14]. Additionally, through direct interaction with the rDNA epigenetics controller, Sir2P, budding yeast Top1 gets recruited to the ribosomal genes wherein it plays a pivotal role in rDNA silencing process [15]. Direct physical interaction between RNA polymerases and DNA topoisomerases has been shown to contribute positively in regulating transcription in multiple systems ranging from bacteria to humans. The β′ subunit of bacterial RNA polymerase was shown to physically interact with the C-terminal domain of bacterial topoisomerase I [16]. Unlike bacteria harbouring a single RNA polymerase, eukaryotic RNA polymerases exhibit division of labour wherein RNA polymerase I transcribes rRNA. RNA polymerase II transcribes mRNA and RNA polymerase III transcribes tRNA, 5srRNA and snRNA. Eukaryotic RNA pol II-Top1 interaction has been explored in *Drosophila* and human systems [17]. During active transcription, direct interaction with RNA pol II helps to recruit Top1 as well as stimulate the latter’s relaxation activity, which in turn facilitates transcription by regulating DNA topology [18]. Bromodomain protein BRD2 in human system interacts, stimulate Top1 activity and help during transcription by suppressing R-loop generation [19]. The proto-oncogene *myc* interacts and facilitates Top1 function at transcriptional sites by forming the ‘topoisome’ complex in humans [20]. Although RNA pol II-Top1 connection is well established, the physical link between Top1 and other two RNA polymerases remain underexplored. Few studies have indicated a genetic and functional interaction between Rpa34 subunit of RNA Pol I and yeast Top1 [21], [22], [23]. On the other hand, limited information is available on RNA pol III-Top1 interaction. Human DNA topoisomerase I was reported to promote RNA Polymerase III mediated transcription reinitiation and termination processes [24]. In budding yeast, two independent high throughput studies based on affinity chromatography followed by mass-spectrometric analyses reported a physical interaction between Topoisomerase I and Rpc82 subunit of RNA Polymerase III [25], [26]. However, detailed characterization of the yTop1-Rpc82 interaction has remained elusive till date. Structurally, Yeast RNA polymerase III comprises seventeen subunits, including five shared across all three RNA polymerases [27], [28], [29], [30], [31], [32]. Among its unique components, the Rpc37–Rpc53 heterodimer facilitates transcription termination [33], [34] while Rpc82 forms a heterotrimer with Rpc34 and Rpc31 that participates in the pre-initiation complex. This Rpc82–34–31 module aids in DNA unwinding to form the open complex [32], [35], [36], [37], [38]. It also helps in stabilizing the transcription bubble in association with Rpc34 [39].

Understanding protein–protein interactions (PPIs) is essential for deciphering cellular functions and regulatory mechanisms. Recent advances in bioinformatics have significantly enhanced the accuracy and speed of structural and interactome predictions, enabling rapid *in silico* analysis. Key applications include protein structure prediction [40], [41], [42], protein structure validation [43], [44], protein-ligand docking [45], [46], [47], protein-protein docking [48], [49], [50], [51], binding free energy estimation [52] and mutational analysis for functional characterization of protein(s) [53], [54], [55]. In this study, we investigated the molecular interaction between yeast DNA topoisomerase I (yTop1) and RNA polymerase III (Pol III) using computational approaches. Due to the lack of a full-length 3D structure of yTop1, we first constructed a model via template-based modeling and validated it through molecular dynamics simulations. This model was then used to explore its interaction with Rpc82, a key subunit of RNA pol III, that is involved in transcription initiation. Using combinatorial protein–protein docking approach, we modeled the yTop1–Rpc82 complex and selected the most favorable binding pose. Molecular dynamics simulations and binding free energy calculations confirmed the stability and strength of the interaction. Comprehensive bioinformatic analysis revealed dynamic features of the interface, offering insights into how yTop1 engages Rpc82 within the larger Pol III holoenzyme. These findings shed light on the molecular basis of Top1–Pol III interaction in budding yeast and lay the groundwork for future functional studies. Given the evolutionary conservation of both proteins, this interaction is likely conserved in humans. In that case, the Top1–RNA Pol III interface in humans may represent a novel target for modulating Pol III activity in diseases such as cancer, where Pol III transcription is often dysregulated[56]. Evaluating druggability of this crucial PPI may open new avenues for targeted cancer therapeutics.

## 2. MATERIALS AND METHODS

### 2.1. Identification of potential Top1 interactors

An initial list of over twenty putative interacting proteins of yeast Topoisomerase I (yTop1) was compiled based on information available on physical interaction from multiple protein–protein interaction databases and literature sources, including the Saccharomyces Genome Database[57], BioGRID [58], STRING [59] and PubMed. To identify transcription-related interactors, specific criteria wer employed such as nuclear localization, direct involvement in transcriptional machinery, and availability of 3D structural information for the candidate protein partner(s). Based on these filters, the Rpc82 subunit of RNA polymerase III was selected, supported by evidence from two independent high-throughput studies employing affinity chromatography followed by mass spectrometry [25], [26].

### 2.2. Template-based modeling of full-length yeast Top1

The full-length amino acid sequence of yeast Topoisomerase I (yTop1) was retrieved from the Saccharomyces Genome Database (UniProt ID: P04786) and analyzed using the InterPro server [60] to identify domain-specific secondary structural elements. Based on this analysis, yTop1 was divided into five segments namely the N-terminal domain (D1, residues 1–142), DNA-binding domain (D2, residues 143–363), core domain (D3, residues 364–589), linker region (D4, residues 590–698), and C-terminal catalytic domain (D5, residues 699–769). Each segment was then subjected to molecular modelling individually, using multiple protein model prediction programs such as RoseTTAFold [61], I-TASSER [62], [63], TrRosetta [64], Ab-initio [65], [66], and DeepFold [67] methods. Few available servers were unable to accept sequences more than 200 residues longs. Due to server limitations on sequence length, segments D1, D3, D4, and D5 were modeled individually using at least three structure prediction tools. D2 was excluded from modeling exercise as its crystal structure is available (PDB ID: 1OIS) [68]. For each segment (D1, D3, D4 and D5), the top five models from each tool were selected based on server-specific scoring criteria. These models were then considered for further quality assessment using the SAVES server, considering ERRAT scores [69], Verify3D parameters [44], Ramachandran plot of stereochemical acceptance and high DOPE score with GA341 score of 1 (Supplementary Table 1).

Based upon the assessment of these scoring parameters, the best model for each yTop1 segment was selected and assembled into a full-length structure using MODELLER[70], incorporating the crystallographically resolved D2 domain (PDB ID: 1OIS). The best model for assembled full-length yTop1 was selected from all the predicted structures based on the validation criteria of ERRAT score, Verify3D, and Ramachandran favoured stereochemical [70, 71], [72] plot utilizing the SAVES server [73]. The structural quality was then refined by energy minimization of the selected assembled model for full-length yTop1. Structural studies have shown that despite sequence divergence, Type IB topoisomerases share a conserved core fold and catalytic mechanism[74], [75]. To benchmark our model, it was compared with the human Topoisomerase I (hTop1, PDB ID: 1K4T), a structurally and functionally conserved orthologue. Sequence identity was checked between yeast and human Topoisomerase I (PDB ID 1K4T), followed by the superimposition of the predicted yTop1 model on hTop1 model to assess the structural comparability of yTop1 segments/domains that share higher sequence identity with their human counterparts.

### 2.3. Molecular docking of yTop1 and Rpc82

To investigate the interaction surface between yTop1 and Rpc82, two molecular docking strategies were employed. Initially, blind docking was performed using the full-length yTop1 model and Rpc82. Given Rpc82’s known interactions with Rpc34 and Rpc160 within the Pol III holoenzyme complex, a targeted docking was then conducted using individual yTop1 domains (D1–D5), with Rpc34/Rpc160-binding residues masked. Comparative analysis of both approaches was carried out identify the most plausible interaction interface between yTop1 and Rpc82.

To investigate the interaction between full-length yTop1 and Rpc82, In order to dock full-length yTop1 with Rpc82, the predicted yTop1 model and the available Rpc82 structure (PDB ID: 6EU0) [76] were used. Blind docking was performed using ClusPro [51], HDOCK [49], HADDOCK[77] servers to identify the possible interaction surface(s) for both the proteins. The top ten docking models from each server were selected based on server-specific criteria.

Binding energies and interaction interfaces were analyzed using PDBePISA[78] and visualized by Biovia Discovery Studio 2021 free version. For each docking model, the percentage contribution of each domain of Rpc82 to the total number of intermolecular non-covalent interactions with yTop1 was calculated and compared across poses to pinpoint the most probable interaction surface on Rpc82 protein. A similar analysis was conducted to identify the Rpc82 binding interface on yTop1 as well. The targeted docking of yTop1 domains at accessible regions of Rpc82 was performed using Cluspro [51] and HDOCK [79] servers followed by binding energy estimation via PRODIGY [80]. Interaction profiling and visualization were again carried out using PDBePISA and Biovia Discovery Studio..

### 2.4. Comparative analysis of yTop1-Rpc82 binding interactions

To further assess the quality of the selected yTop1–Rpc82 docking model, the ratio of binding energy to interaction surface area was evaluated and compared with other established heterodimeric protein-protein complexes having similar ranges of binding energies and surface areas, following previously reported method [81]. The docking results of the yTop1-Rpc82 complex were evaluated and validated by comparing their interaction characteristics against heterodimeric protein complexes of similar size and interaction profiles. Interaction surfaces were calculated using the EMBL PDBsum program[82], while binding energy (ΔG) values were estimated using the PRODIGY server [80]. This comparative analysis was performed to validate the docking model by benchmarking its interaction characteristics against known protein-protein complex standards.

### 2.5. Molecular dynamics simulations

The predicted yTop1 model was subjected to molecular dynamics (MD) simulation analysis using GROMACS [83], [84] version 2024. AMBER99SB protein force field [85] was utilised to define the topology file. The protein-in-water complex was accommodated in a box of 2081.95 nm^3^ cube. Charge neutralization was achieved by the application of 12 Cl^-^ ions. Steepest descent [86] and conjugant gradient algorithms [87] were used for energy minimization followed by the system equilibration in the NVT and NPT ensemble at 300 K for 0.1 ps time constant. Final MD run was performed with a step size of 2 fs. Stability and dynamics of the protein were assessed by calculating different parameters like RMSD, RMSF, Surface Accessible Surface Area (SASA) etc. of simulation trajectory.

Similarly, to assess the structural dynamics and stability of the selected Top1-Rpc82 docking complex, molecular dynamics (MD) simulation studies were conducted for 100ns using GROMACS platform [83], [88]. AMBER99SB protein force field [89] was applied to perform these studies. The protein in water complex was measured to be occupying a cubic volume of 5505.45 nm^3^, which made neutral by applying 24 Cl ions. Energy minimization of the system was performed by Steepest descent [86] and conjugant gradient algorithms [87], followed by the system equilibration in the NVT and NPT ensemble at 300 K at 0.1ps time constant. The final MD run followed the 2fs step size to attain the total of 100 ns simulation. PDBePISA server [78] and Biovia Discovery studio software [90] was employed to sense the interacting residues of the two proteins during the simulation course by choosing different coordinate frames of regular interval. Major interacting zone during the simulation was determined and its interaction dynamics was assessed by calculating the RMSD, RMSF, Rg and SASA values of the interacting residues/surface followed by extrapolation of the values on the respective graphs of the whole complex.

### 2.6. MM-PBSA analysis of yTop1-Rpc82 complex stability

Protein-protein interaction energetics were quantified using the Molecular Mechanics Poisson-Boltzmann and Surface Area (MM-PBSA) methodology [52], a computational approach for precise binding free energy determination through molecular dynamics (MD) simulations. The binding free energy was calculated using the fundamental thermodynamic equation

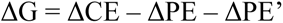

where ΔCE represents the complex free energy, and ΔPE and ΔPE’ denote individual protein free energies. The comprehensive MM-PBSA binding free energy equation,

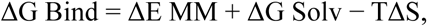

systematically decomposes molecular interaction energetics into molecular mechanics energy (ΔE MM), solvation free energy (ΔG Solv), and entropic contributions. Molecular mechanics energy was dissected into bonded (ΔE bonded) and non-bonded (ΔE nonbonded) components, further resolved into van der Waals (ΔE vdW) and electrostatic (ΔE elec) interactions. Solvation free energy was analysed through polar (ΔG PB) and non-polar (ΔG SA) contributions. Molecular dynamics simulations were performed using GROMACS with a 100 ns trajectory, generating 200 snapshots at 500 ps intervals under standardized conditions: 0.154 M salt concentration, with dielectric constants of 1.0 (internal) and 80.0 (external). The gmx MM-PBSA [52] computational interface enabled precise binding free energy calculations, incorporating an interaction entropy (IE) module [91] to directly quantify entropic contributions from MD simulation ensembles.

The final binding free energy determination followed the equation

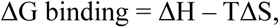

where ΔG binding represents Gibbs free energy of protein-protein interactions, ΔH accounts for molecular interaction enthalpy, and TΔS captures conformational entropy. This methodology provides a rigorous computational framework for investigating protein-protein interaction energetics at atomic resolution, combining trajectory generation, snapshot analysis, energy component decomposition, and entropic contribution assessment. While offering detailed molecular insights, the approach necessitates complementary experimental validation to fully elucidate complex intermolecular interactions.

## 3. RESULTS

### 3.1. Structural Modeling of Full-Length Topoisomerase I from Budding Yeast

Accurate three-dimensional (3D) structural information is essential for *in-silico* molecular docking studies. While the structure of Rpc82 is available in the RCSB-PDB (PDB ID: 6EU0), yeast Topoisomerase I (yTop1), a 769-residue protein, lacks a complete structural model. The only crystal structure available (PDB ID: 1OIS) covers the DNA-binding domain (residues 141–363), leaving the rest of the protein unresolved. Even the AlphaFold prediction (AF-P04786-F1) omits the N-terminal domain (NTD), which is known to mediate interactions with several cellular proteins such as Rpb1, Sir2, and Tof1. Thus, it was necessary to build a model for full-length yTop1. Initial homology modeling using SwissModel program [41] based on 51.69% sequence identity with the human orthologue, failed to capture the first 128 N-terminal residues and a significant C-terminal stretch. Therefore, to enable reliable interaction studies, an alternative modeling strategy was adopted to generate structural information for key regions of yTop1, including the NTD (D1) and linker region (D4), which were missing from existing AlphaFold and homology-based models.

To ensure unbiased 3D structure prediction, each yTop1 domain (D1, D3, D4, D5) was modeled using multiple programs and validated using previously described rigorous criteria. All selected models showed high structural accuracy, with ERRAT scores above 95% and Verify3D scores exceeding 85%, indicating strong sequence–structure compatibility.

Ramachandran plot analysis revealed over 91% residues in favored regions with minimal presence in disallowed zones (Supplementary Figure 1, Supplementary Table 1). The validated domain models were then assembled using MODELLER to generate the full-length yTop1 structure (Figure 1). The final model was selected based on optimal validation metrics such as am ERRAT score of 93.43%, Verify3D score of 86.35%, 94.3% residues in favored Ramachandran regions, and only 0.1% in disallowed regions. Additional parameters such as GA341 score of 1, DOPE score of −79,285.81, and molPdf score of 4632.84 further confirmed the structural integrity and successful assembly of individual domains into a reliable full-length structure.

**Figure 1.**
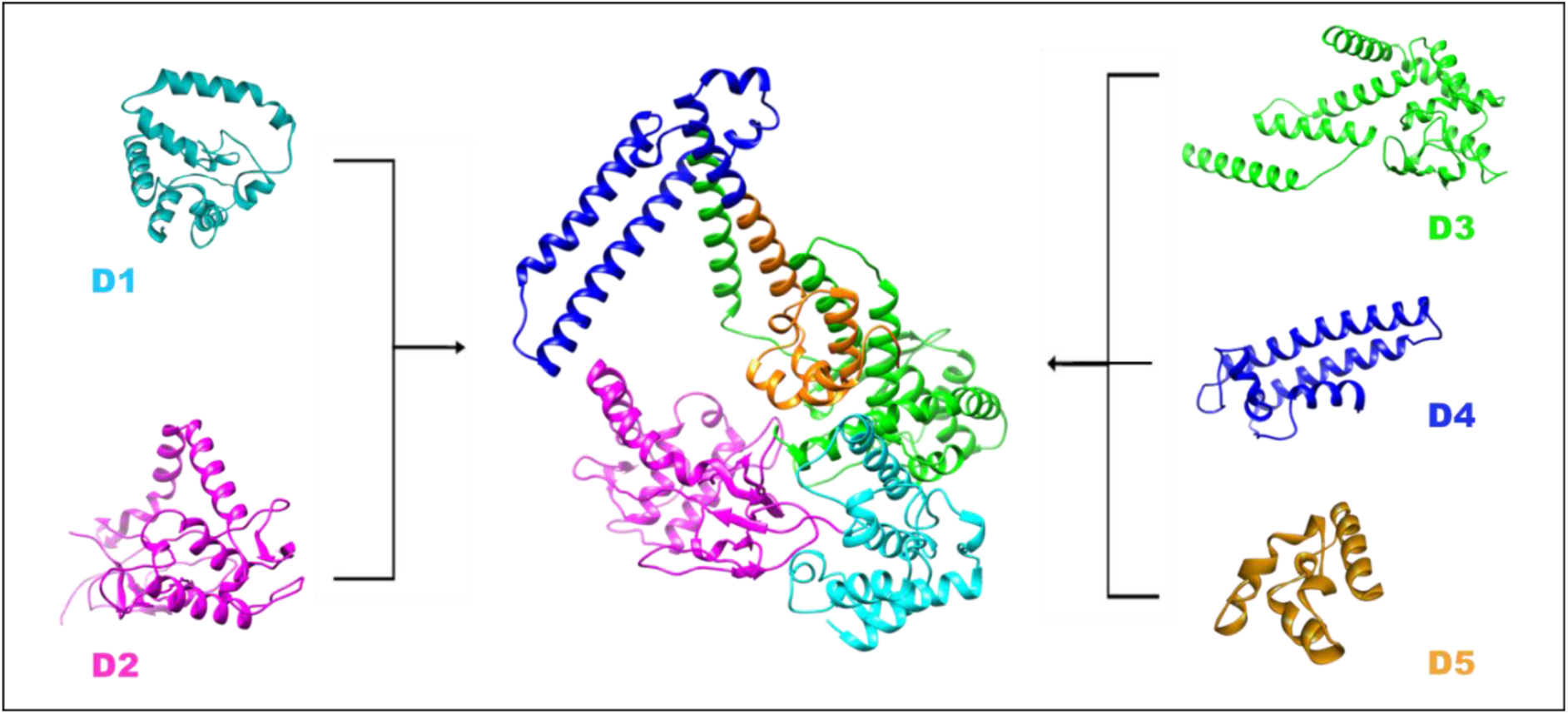
Domain-wise representation of modeled Topoisomerase I from budding yeast. The full-length yeast Topoisomerase I (yTop1) structure was assembled by integrating predicted models of domains D1, D3, D4, and D5 with the experimentally resolved D2 domain (PDB ID: 1OIS). The central 5-colored image illustrates the assembled model, with each domain distinctly color-coded, D1 (cyan), D2 (purple), D3 (green), D4 (blue), and D5 (orange), highlighting their spatial arrangement within the complete protein structure.

### 3.2. Comparative assessment of yTop1 model against human Topoisomerase I crystal structure

The UniProt entry for yeast Topoisomerase I (yTop1, P04786) reveals conservation of key catalytic residues and domain architecture with its human counterpart. Sequence similarity analysis identified human Topoisomerase I (hTop1, PDB ID: 1K4T) as the closest homolog, with highest similarity in the D2 and D3 regions. Overall, yTop1 shares 51.69% sequence identity and 45% similarity with hTop1. To ascertain whether our predicted model for yTop1 model successfully captures the essential structural features of the enzyme, especially around the active-site tyrosine and DNA-binding motifs, structural comparison of our predicted model was performed with hTop1 structure. Since the available structural model of the human Topoisomerase I (hTop1) protein lacks information on the N-terminal 173 residues, thus, D1 of yTop1 could not be compared with hTop1 structure. Superimposition using RMSD (Root Mean Square Deviation) is a widely accepted method for assessing the quality of a predicted model. When a predicted model is superimposed onto a reference structure derived from crystallographic data, RMSD quantifies the average distance between corresponding atoms after optimal alignment. A lower RMSD (< 2 Å) indicates a closer match and a more accurate model. [92], [93] [94]. The predicted full-length yTop1 model was superimposed onto the hTop1 structure and based on the alignment score, the overall RMSD value was observed to be 1.021 Å (Supplementary Figure 2A). Additionally, superimposition of yTop1 domains D2–D5 onto the AlphaFold-predicted structure (Figure 2A) yielded an RMSD of 0.7 Å. Notably, while the AlphaFold model lacks recognizable secondary structures in D1 region except for a single α-helix (residues 105–125), our predicted full-length model successfully recapitulates this feature (Figure 2B), further validating its accuracy.

**Figure 2.**
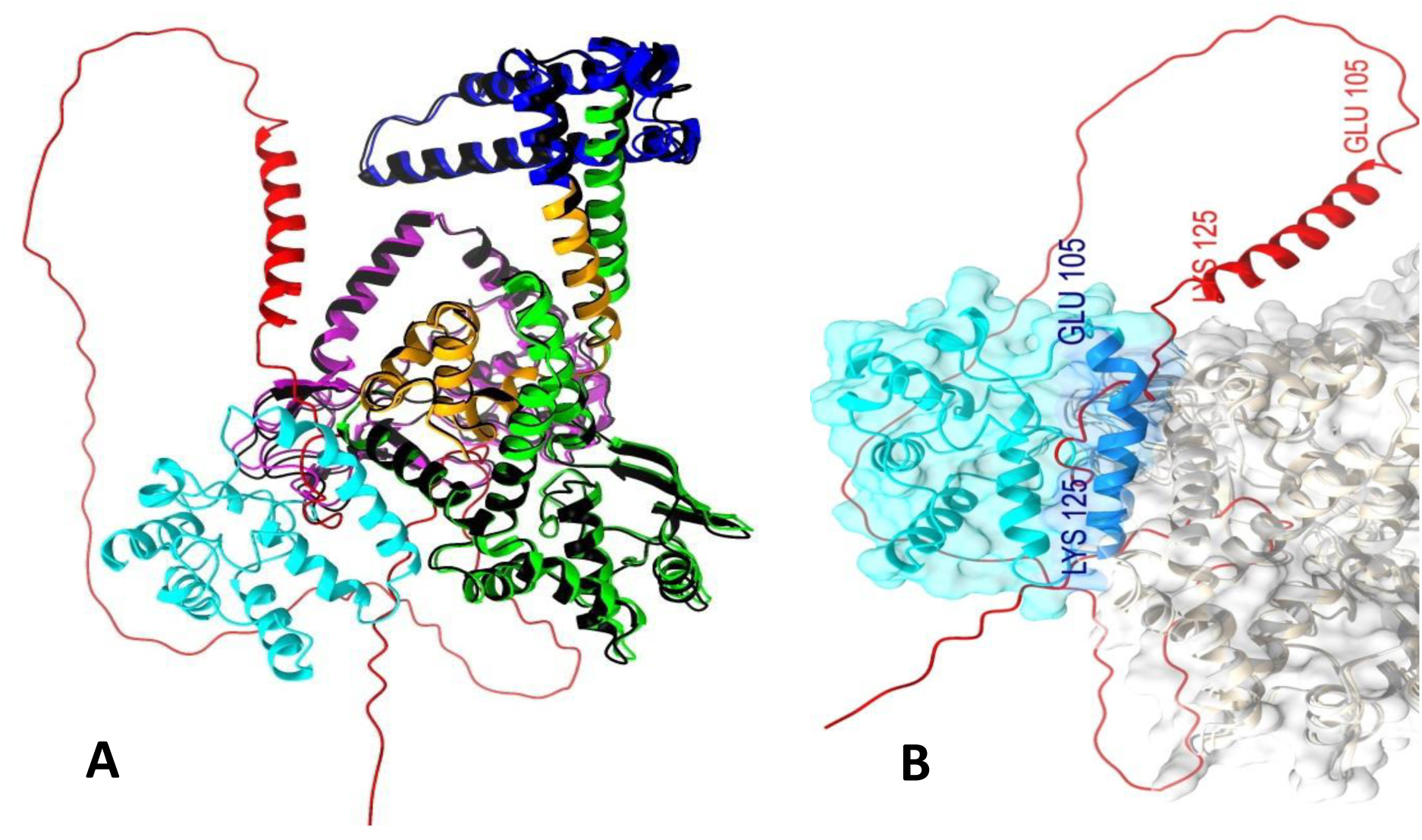
Structural comparison of the predicted full-length yTop1 model with AlphaFold model of yTop1. (A) Superimposition of the predicted yTop1 model (D1–cyan, D2–magenta, D3–green, D4–blue, D5–orange) onto the AlphaFold-predicted structure (D1–red and dark grey for rest of the length) shows negligible RMSD across most regions, except for D1 due to the lack of defined secondary and/or tertiary structure in the AlphaFold model. (B) Enlarged view of the D1 region (cyan) from the predicted model, highlighting the successfully resolved α-helix (residues 105–125, dark blue), which aligns with the only recognizable secondary structure in the D1 domain of AlphaFold model (same helix shown in red).

To further evaluate the individual domain models of yTop1, sequence and structural comparisons were performed against the human Topoisomerase I (hTop1). The D2 domain of yTop1 shares 51.14% sequence identity and 46% similarity with residues 213–431 of hTop1, yielding an RMSD of 0.941 Å upon superimposition (Supplementary Figure 2B). Domains D3 and D5 showed 54.02% and 54.9% sequence identity, and 44% and 49% similarity with hTop1 residues 432–663 and 695–765, respectively. Their RMSD values were 0.805 Å (Supplementary Figure 2C) and 1.29 Å (Supplementary Figure 2D), indicating strong structural alignment. Overall, RMSD fluctuations for D2, D3, and D5 remained below 2 Å, confirming that the predicted *Saccharomyces cerevisiae* Top1 model aligns very closely with the human Top1 reference, particularly in the conserved regions. In contrast, the D4 linker region lacked sequence identity with any resolved eukaryotic topoisomerase structures available in the RCSB-PDB, suggesting it may be unique to yeast or highly divergent. Given that linker regions in Topoisomerase I are typically flexible and involved in transient or condition-specific interactions, structural comparison for D4 domain was conducted using the AlphaFold-predicted model of yTop1. The resulting structural deviation (RMSD) of 1.253 Å supports the structural reliability of our predicted model (Figure 2 and Supplementary Figure 3).

### 3.3. Molecular simulations to assess Structural dynamics of the yTop1 model

Molecular Dynamics (MD) simulations offer valuable insights into protein stability, flexibility, and conformational dynamics. To validate the predicted yTop1 model, MD simulation was performed. As shown in Figure 3A, RMSD values stabilized around 0.7 nm after 45 ns and decreased below 0.6 nm after 65 ns, indicating structural stability. RMSF analysis of all amino acid (aa) residues of the yTop1 (Figure 3B) revealed higher fluctuations in the N-terminal domain (D1) and linker region (D4), ranging from 0.6–0.7 nm, while D2, D3, and D5 domains showed lower fluctuations (0.2–0.4 nm), suggesting greater rigidity. The residues of the NTD and linker region exhibited a higher level of fluctuations and thus have higher flexibilities than the remaining three domains. The radius of gyration (Rg) remained around 3.55 nm (Figure 3D), with a slight decreasing trend after 60 ns, indicating compactness. SASA values were consistently ∼440 nm² throughout the simulation (Figure 3C), reflecting stable surface exposure. Collectively, these results confirm the structural integrity and suitability of the predicted yTop1 model for downstream protein–protein docking studies.

**Figure 3:**
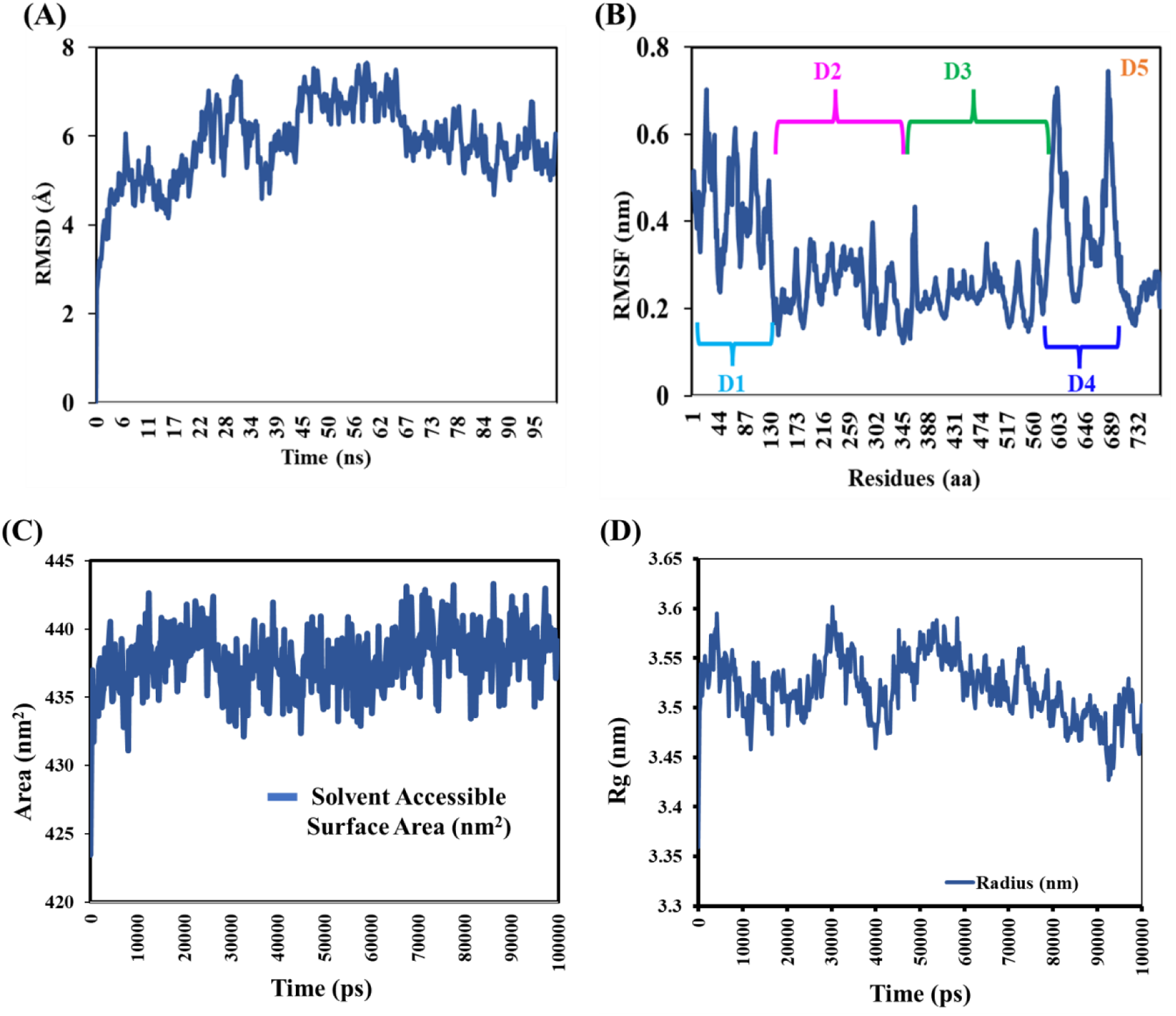
Molecular Dynamics Simulation Analysis of Predicted yTop1 Model (A) RMSD plot showing structural stability of the full-length yTop1 model, with deviations consistently below 0.8 nm throughout the simulation. (B) RMSF analysis indicating higher flexibility in domains D1 and D4 compared to the more stable D2, D3, and D5 regions. (C) Solvent Accessible Surface Area (SASA) profile reflecting dynamic surface exposure during simulation. (D) Radius of gyration plot demonstrating compactness and overall structural integrity of the predicted model.

### 3.4. Molecular docking and interaction analysis between yTop1 and Rpc82

Rpc82 is the third largest and functionally important subunit of the RNA polymerase III (Pol III) complex in *Saccharomyces cerevisiae*, which plays a pivotal role in the transcription of small non-coding RNAs, including tRNAs and 5S rRNA [76]. Structurally, Rpc82 consists of four winged-helix domains (WH1–WH4) and a C-terminal coiled-coil (CC) domain. It anchors to the clamp domain of the Pol III cleft and interacts with duplex DNA downstream of the transcription bubble, contributing to transcription complex stability [95]. A disordered insertion within its WH3 domain facilitates interaction with Brf1, a TFIIB-related transcription factor, during pre-initiation complex formation [95], positioning Rpc82 as a key mediator of protein–protein contacts within the Pol III pre-initiation complex. Given its strategic location within Pol III holoenzyme and evidence from two independent affinity chromatography studies, Rpc82 is proposed to serve as a critical interface for binding by regulatory proteins like Topoisomerase I, especially during transcription of tRNA genes wherein relieving torsional stress is essential.

In order to explore the molecular interactions between Rpc82 and yTop1, molecular docking was performed initially with full-length predicted yTop1 and full-length Rpc82. Two distinct docking interfaces for yTop1 were identified on the Rpc82 surface, with binding energy details summarized in Supplementary Table 2. Among these, the most favorable docking pose, characterized by a strong negative ΔG value, highlighted the WH2–WH3 region of Rpc82 as the primary interaction surface (Figure-4A, 4B and Supplementary Figure 5). This interface was prioritized for further analysis based on both energetic and structural considerations. Unlike the WH1–WH4 and CC domains, which are structurally committed to stable interactions within the Pol III core (notably with Rpc160 and Rpc34, as seen in the RNA Pol III holoenzyme structure (PDB ID: 6EU0) [96], [97], the WH2–WH3 region remains spatially accessible (Supplementary Figure 4). Its peripheral location and involvement in transient interactions, such as with the Brf1 during pre-initiation complex formation, make it a plausible docking site for regulatory proteins like Topoisomerase I. Moreover, selecting an interface outside the core-engaged domains avoids steric clashes and preserves the integrity of the Pol III assembly. Therefore, the best docking pose involving accessible regions i.e. the WH2–WH3 surface (Supplementary Figure 5) was considered for further evaluation. Overall, the WH2–WH3 region represents a structurally viable and biologically relevant surface for Top1 interaction, aligning with both docking energetics and known functional roles of Rpc82. The interaction surface on Topoisomerase I, as illustrated in Figures 4C, 4D, and 4E, highlights the involvement of discrete regions within its D1 and D3 domains primarily.

**Figure 4:**
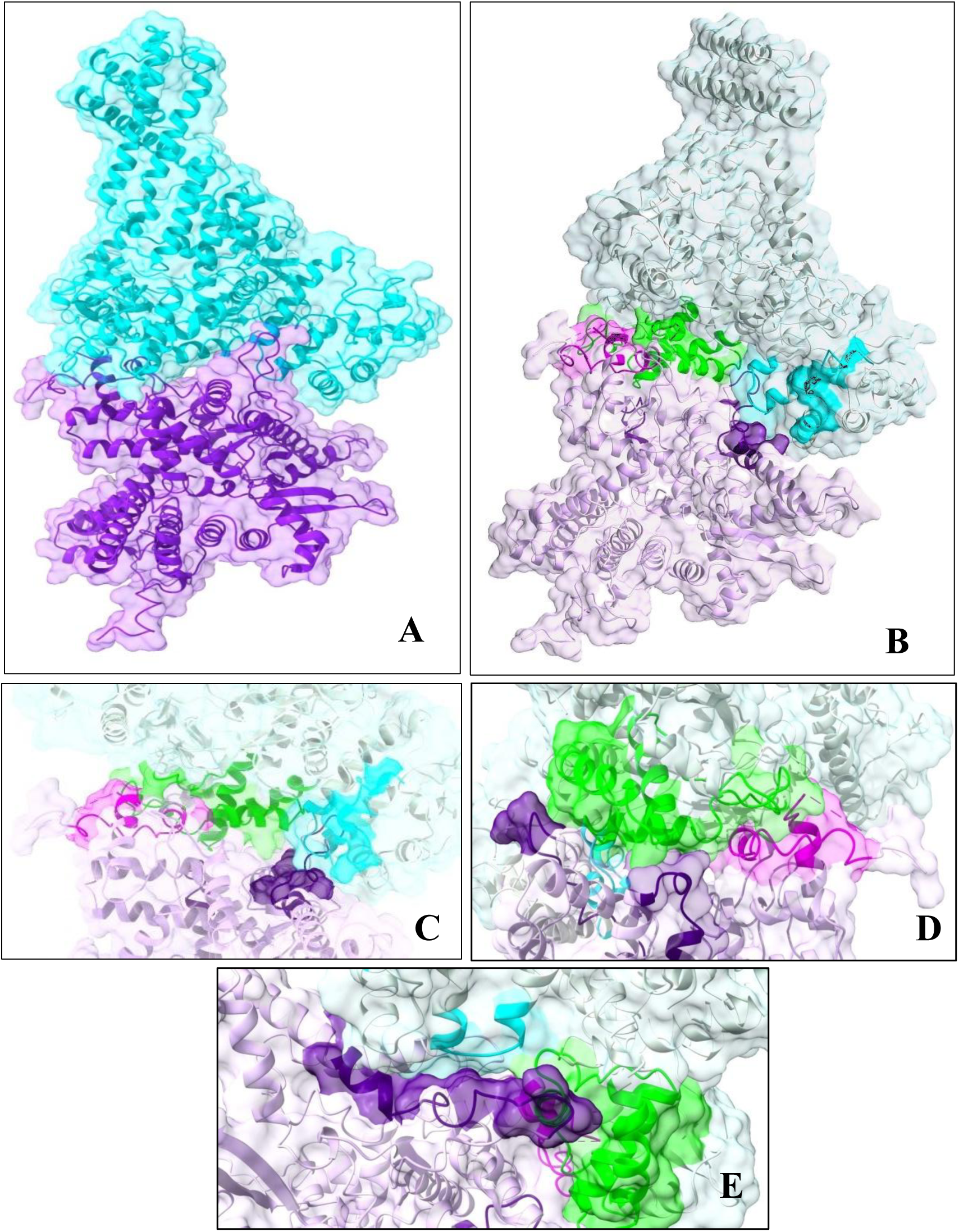
(A) Selected docking model of *S. cerevisiae* Topoisomerase I (yTop1, cyan) in complex with Rpc82 (violet). (B) Overview of the full yTop1–Rpc82 docking complex, with non-interacting regions shown in grey and interaction surfaces highlighted in color. (C–E) Close-up views of the interaction interface: D1 (cyan) and D3 (green) domains of yTop1 engage with the WH2 (violet) and WH3 (purple) domains of Rpc82. Non-interacting surfaces of yTop1 and Rpc82 are shown in faint blue and pale violet, respectively.

To further strengthen our findings and capture the most plausible interaction surface on Rpc82, a targeted docking approach was employed using individual domains of yTop1 and the accessible regions of Rpc82. Top-scoring docking models were selected based on binding free energy calculations from the PRODIGY server [80]. The analysis **(supplementary Table 3**) revealed that yTop1 domains D1, D2, D4, and D5 interact with Rpc82’s WH2–WH3 region (residues 140–196 and 272–369), while the D3 domain also engages this surface and forms additional contacts with the WH1 domain. Among all, D1 and D3 showed the strongest binding affinities, indicating their central role in the interaction. Moreover, the convergence of both blind docking (full-length yTop1) and targeted docking using individual domains of yTop1 on the WH2–WH3 region of Rpc82 protein supports its relevance as the primary binding interface. Furthermore, the WH3 domain’s known interaction with Brf1 highlights its *in vivo* accessibility and functional flexibility, even within the Pol III holoenzyme context [95].

### 3.5. Binding free energy analysis of yTop1-Rpc82 complex

As outlined in the methods section, molecular docking of full-length yTop1 with Rpc82 was performed using multiple servers. From each docking solution, the top ten scoring complexes were analyzed to assess binding energies, interaction frequencies, and domain-specific contributions of Rpc82 towards complex stability. Detailed interaction analysis for the top scoring models is provided in Supplementary table 2. The selected docking model as presented in Figure 4A revealed that yTop1 predominantly interacts with the WH2 and WH3 domains of Rpc82. This model exhibited a favorable binding energy of –11.5 kcal/mol as depicted in Table 1. This table presents the total number of intermolecular contacts between yTop1 and Rpc82 in the selected complex. The relative contribution of each Rpc82 domain to the interaction interface was quantified by calculating the percentage of contacts formed, offering insights into the involvement of WH2 and WH3 domain in stabilizing the yTop1–Rpc82 interaction.

**Table 1:** Domain-wise interaction analysis of Rpc82 with the full-length yTop1 for the selected docking complex of yTop1-Rpc82.

| Total number of interaction in TopoI-Rpc82 interface | Rpc82 domain-specific contribution to the overall interaction | | Top I domain-specific contribution to the overall interaction | | $\Delta G$ |
| --- | --- | --- | --- | --- | --- |
|  | WH2 | WH3 | D1 | D3 |  |
| 39 | 64.1% | 35.9% | 13% | 87% | -11.5 |

Total number of interactions were found to be 39 between the two proteins as mentioned in the first column. Interaction of yTop1 at different domains of Rpc82 surface was estimated and converted to percentage values to determine the contribution of each of the two Rpc82 domains (WH2 and WH3) towards the total binding interactions observed with yTop1.

As mentioned previously, the second predicted binding surface on Rpc82 (formed by WH1-WH4, CC domain) was excluded from further analysis due to its involvement in RNA Pol III holoenzyme assembly, where interaction with core subunits would likely cause steric hindrance. Heatmap analysis (Supplementary table 4) revealed domain-wise energy contributions and interaction percentages across Rpc82. However, since the WH1–WH4 and coiled-coil (CC) regions remain pre-occupied by Rpc160 and Rpc34 within the RNA Polymerase III holoenzyme (Supplementary Figure 4), the WH2–WH3 domains of Rpc82 emerged as the only accessible and energetically favorable interface for Topoisomerase I binding. From the perspective of yTop1, amino acid residues within the D1 and D3 domains primarily and few residues from the D2 domain contribute to the formation of a potential interaction surface on Topoisomerase I (Figure 4B-E).

### 3.6. Validation of yTop1-Rpc82 complex through comparative structural analysis

As described previously, the docking model in which yTop1 preferentially interacts with the WH2–WH3 domains of Rpc82 was selected for further analysis due to its biological plausibility. To evaluate the accuracy and reliability of this complex, its protein–protein interaction energy and surface area were compared against a benchmark dataset of heterodimeric protein complexes from the RCSB-PDB (Supplementary Table 5). As shown in Figure 5A, the energy plot of the selected yTop1-Rpc82 docking complex demonstrated that the binding energy values of yTop1-Rpc82 complex falls within the range observed for natural heterodimeric protein-protein interactions, indicating a stable and favorable interface. These results indicate that the selected yTop1-Rpc82 complex exhibits a stable and energetically favourable interaction. Similarly, the whisker plot of the energy/surface area values (Figure 5B) of the yTop1-Rpc82 complex was placed below −0.008, which confirmed the interaction efficiency, represented by the energy-to-surface-area ratio. The obtained value for our complex aligns closely with the standard ranges between −0.014 to −0.005. The calculated energy-to-surface-area ratio for the yTop1–Rpc82 docking complex closely aligns with benchmark values derived from well-characterized heterodimeric protein interactions. This clustering within the biologically accepted range reinforces the notion that the predicted complex mimics natural protein–protein interfaces and further strengthens the structural reliability of the docking model and its biological plausibility.

**Figure 5:**
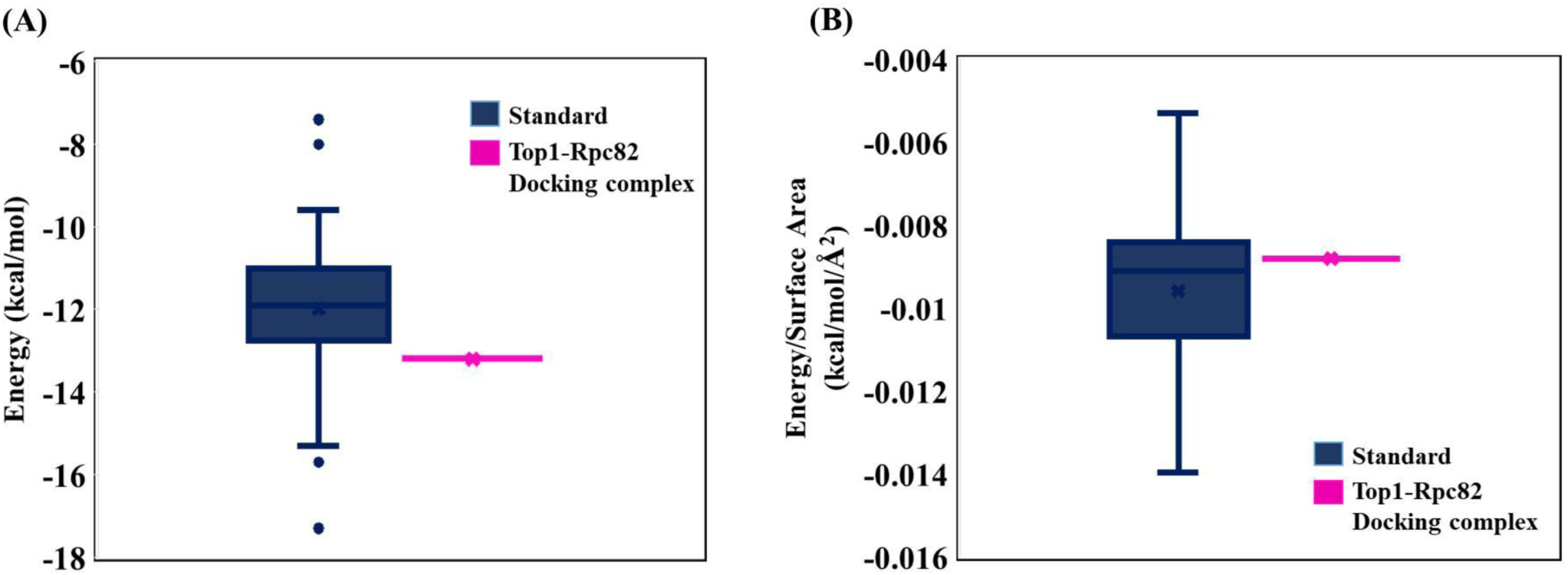
Comparative analysis of yTop1-Rpc82 docking complex against the reference dataset of experimentally validated heterodimeric protein complexes (A) Binding energy plot comparing the yTop1–Rpc82 docking complex with a reference dataset of experimentally validated heterodimeric protein complexes. The selected model shows energy values within the typical range observed for stable protein–protein interactions. (B) Whisker plot illustrating the energy-to-surface-area ratio of the yTop1–Rpc82 complex relative to benchmark heterodimeric interfaces. The ratio falls within the biologically relevant range (–0.014 to –0.005), supporting the interaction efficiency and structural plausibility of the predicted complex.

### 3.7. Molecular Dynamics simulation validates stability of yTop1–Rpc82 complex

Binding free energy calculations using the gmx_MMPBSA module over a 100 ns simulation confirmed the energetic favorability of the yTop1–Rpc82 complex. A stable binding state was achieved early in the simulation and maintained throughout, with an average ΔG value of – 104.97 kcal/mol. The root mean square deviation (RMSD) analysis (Figure 6A) showed initial deviations between 0.4–0.6 nm, followed by equilibration within the first 10 ns, indicating a moderately stable configuration. Although minor fluctuations occurred after 20 ns, RMSD values remained below 0.7 nm for the entire trajectory. These minimal deviations and stable stable RMSD trajectories of the model suggest that the complex retains structural integrity and dynamic stability under simulated physiological conditions. The predicted docking pose thus emerged as energetically favourable, structurally stable and represents a biologically plausible interaction between yTop1 and Rpc82.

**Figure 6:**
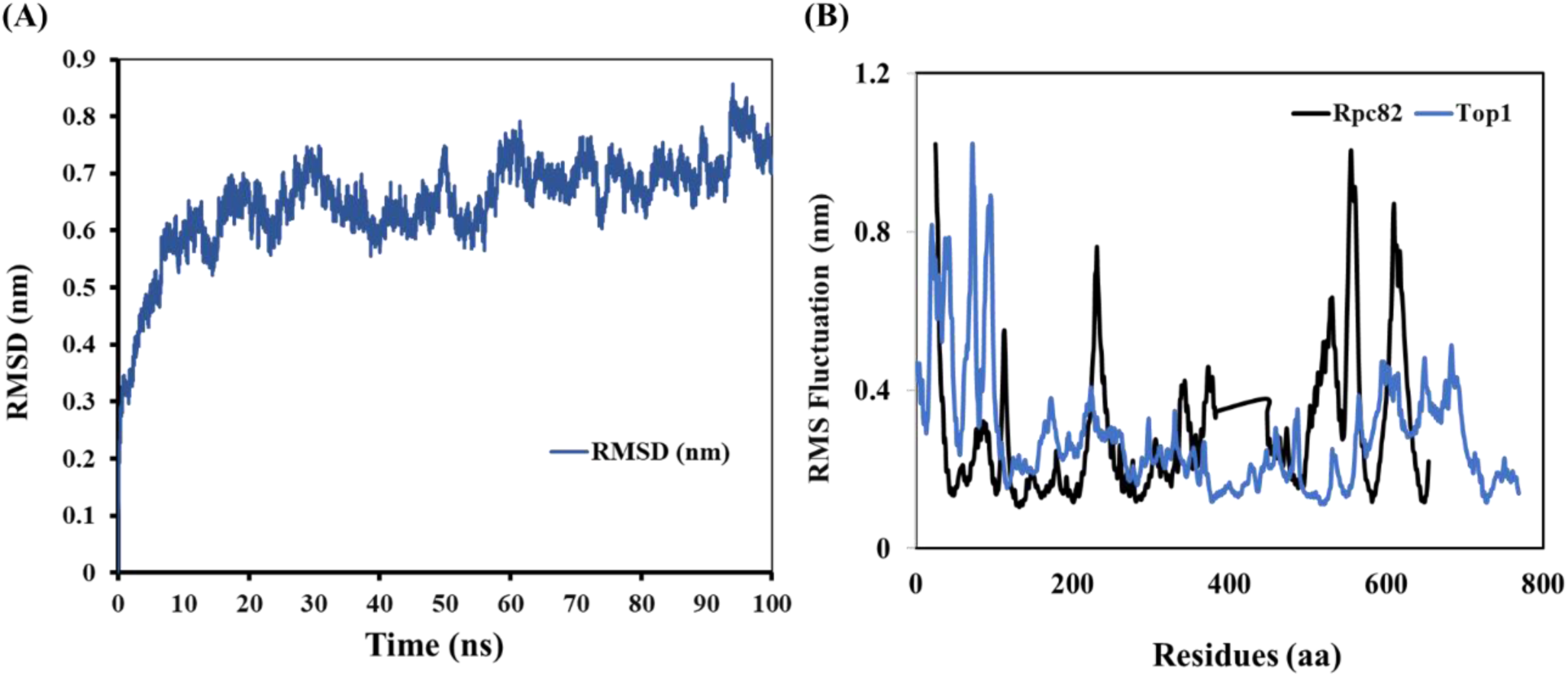
Molecular dynamics simulation confirms structural stability of the yTop1–Rpc82 complex. (A) RMSD plot showing the overall stability of the yTop1–Rpc82 complex throughout the simulation. RMSD values remained consistently below 0.8 nm, indicating minimal structural deviation and a stable interaction. (B) RMSF plot illustrating residue-level flexibility across both proteins. Blue trace represents yTop1 and black trace represents Rpc82, highlighting domain-specific fluctuations and dynamic regions within the protein complex.

Root mean square fluctuation (RMSF) analysis was performed to assess residue-level flexibility within the yTop1–Rpc82 complex. As shown in Figure 6B, residues at the interaction interface exhibited limited fluctuations, with RMSF values below 0.3 nm, indicating high structural stability. Specifically, Rpc82’s WH3 and WH4 domains, along with yTop1’s DNA-binding (D2), core (D3), and C-terminal (D5) domains, demonstrated minimal flexibility and higher stability over the simulation trajectories. In case of Rpc82, amino acid residues belonging to the WH4, cleft loop (aa. 550-563) and coiled-coil (CC) regions showed higher RMSF values. In contrast, residues within NTD (D1) of yTop1 and CC domain of Rpc82 exhibited maximum fluctuations of up to 1.0 nm, reflecting structural flexibilities under simulated physiological conditions. These observations highlight the interplay between stability at the binding interface and intrinsic flexibility in the other regions of the protein-protein complexes under dynamic physiological conditions.

Solvent Accessible Surface Area (SASA) and Radius of Gyration (Rg) data revealed the structural compactness of the whole protein complex as well as the interaction surface. The average SASA values of individual interacting regions were found to be ∼ 320 nm^2^ for yTop1 and ∼280 nm^2^ for Rpc82. These values are much lower than the value observed for the full complex, ∼ 800 nm^2^, and this was maintained till the end of the simulation (Supplementary figure 6). The residue-specific SASA values of the interacting surface were found to be placed at ∼ 1.5 - 2 nm^2^ for both the proteins. The Rg values of the full-length yTop1-Rpc82 simulation suggested that the complex remains stable at ∼ 4.2 nm. The interaction surfaces for yTop1 and Rpc82 showed a similar trend of compactness by ∼ 3.3 nm and ∼ 3.5 nm respectively (Figure 7D).

**Figure 7:**
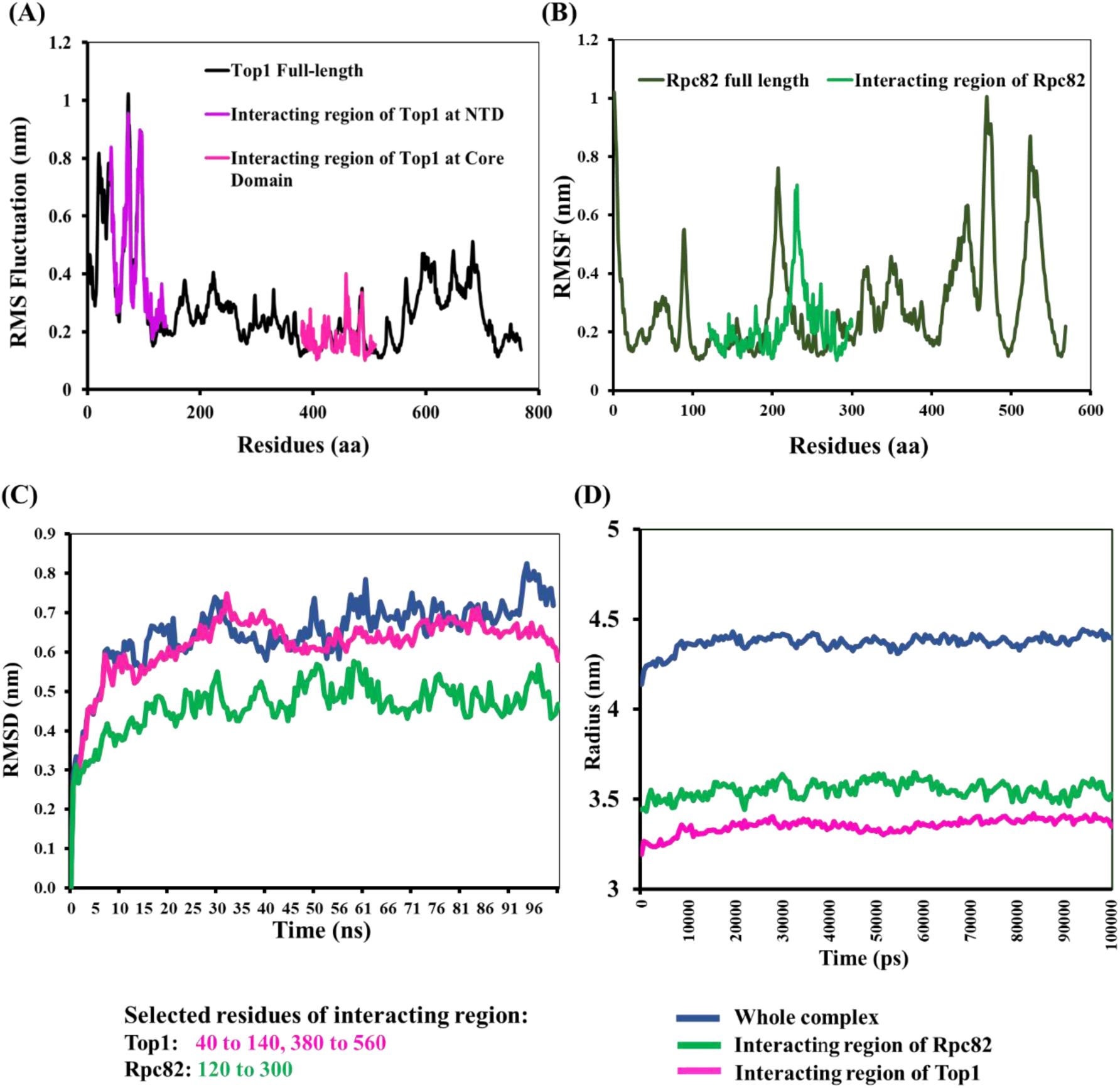
Molecular Dynamics based evaluation of Topoisomerase I (yTop1) and RNA Polymerase III Subunit Rpc82 Interaction, (A) RMSF profile of full-length yTop1 highlighting residue-level fluctuations, particularly at interaction regions. (B) RMSF analysis of Rpc82 showing domain-specific flexibility, with WH3 and WH4 regions exhibiting high stability. (C) RMSD trajectory of the yTop1–Rpc82 complex over 100 ns, indicating overall structural stability with minimal deviation. (D) Radius of gyration (Rg) plot demonstrating compactness and conformational stability of the complex throughout the simulation.

### 3.8. Key interaction hotspots contributing to complex stability during simulation

In-depth analysis of the interaction surface between the two proteins during the simulation course was achieved. Persistent interactions were visualized between the amino acid residues of yTop1 and Rpc82 across the simulation trajectory. Amino acid residues belonging to two separate regions, namely NTD (40 −140 aa of D1 domain) and core domain (380- 560 aa of D3 domain) were observed in forming the interaction surface of yTop1. On the other hand, residues of a single interaction surface (formed by WH2 and WH3 domains) of Rpc82 (residue position: 120-300, 447,448) was revealed by this assessment. Certain residues of yTop1 were revealed to be critical for maintaining this interaction throughout the simulation course, including D50, E126, E127 residues of N-terminal domain (D1) as well as Y385, R395, K400, E424, E427, D493, R494, K502, N506 residues belonging to the core-domain (D3) of the enzyme.

Looking at the interaction surface on Rpc82 protein, several residues namely R145, D148, E149, Y185, V192, T264, D266, T269, S270, R272 of WH2 domain and A272, H448 residues of WH3 domains were found to be significant for maintaining the interaction with yTop1 almost for the entire period of simulation (Supplementary Table 6). Due to very high interaction frequencies observed for the above-mentioned residues of both the proteins, they are considered to be consistently engaged throughout the simulation, thereby indicating formation of a stable protein-protein binding interface. Interestingly, during simulation, specific residues such as K221, K224, R228, K255, L259 of Rpc82 appears to form a transient interacting patch. The consistency of the interaction pattern along with the development of additional interacting patch suggested the ability of the complex to adapt and accommodate itself to maintain interaction by undergoing conformational changes in the dynamic physiological condition.

Consistent stability of the interaction patterns indicated towards the functional relevance of this PPI. For further detailed examination of the interaction surface, SASA (Supplementary Figure 6) and Rg (Figure 7D) of the interacting surface residues of both theproteins were calculated and compared with the whole complex. Residue-wise change in the accessible surface area were also measured for the selected residues of both the protein and the result indicated the sustained anchorage between the two proteins throughout the duration of simulation.

### 3.9. Interfacial dynamics of yTop1-Rpc82 binding

The comprehensive analysis of root-mean-square fluctuation (RMSF) and root-mean-square deviation (RMSD) revealed distinctive dynamic behaviours for yTop1 and RNA polymerase III subunit Rpc82 during molecular dynamics simulations. Figure 7A demonstrates the RMSF profile of yTop1, highlighting significant fluctuations across its full-length structure. The interacting regions at the N-terminal domain (NTD) and core domain (40-140 and 380-560 residues respectively) exhibited notably different fluctuation patterns compared to the full-length protein. Specifically, the interacting regions showed more constrained fluctuations, with peak amplitudes ranging between 0.4-1.0 nm, indicating potential stabilization of these critical interaction sites. Figure 7B illustrates the RMSF characteristics of Rpc82, with the full-length protein and its interacting regions (120-300 residues) displaying distinct fluctuation profiles. The interacting region demonstrated more pronounced structural dynamics, with fluctuation peaks reaching up to 0.8 nm, suggesting potential conformational flexibility critical for protein-protein interactions. The RMSD analysis (Figure 7C) tracked the structural stability of the proteins over the 100 ns simulation time. The whole complex, interacting regions of Rpc82, and interacting regions of yTop1 showed progressively stabilizing behaviour, with RMSD values converging around 0.5-0.7 nm. This output indicates a robust and consistent protein-protein interaction. Figure 7D provides a long-time scale stability analysis, demonstrating the consistent radius of gyration for the whole complex, interacting regions of Rpc82, and yTop1. The relatively stable values (ranging between 3.5-4.5 nm) suggest that the structural integrity of the complex was maintained throughout the extended simulation period.

The overall Gromacs free energy of the selected complex reached the values around - 8310000 kJ/mol (Figure 8A) when passing through the 100 ns of MD simulation trajectories. The MM-PBSA binding free energy of the interacting surface made by the mentioned residues was calculated. The results indicated a strong and stable binding between yTop1 and Rpc82 during the molecular dynamics timeframe. The calculated average ΔG value of the interaction patch (Figure 8B) was −302.56 KJ/mol wherein Van der Walls interactions were found to be a significant contributor along with the electrostatic interactions.

**Figure 8:**
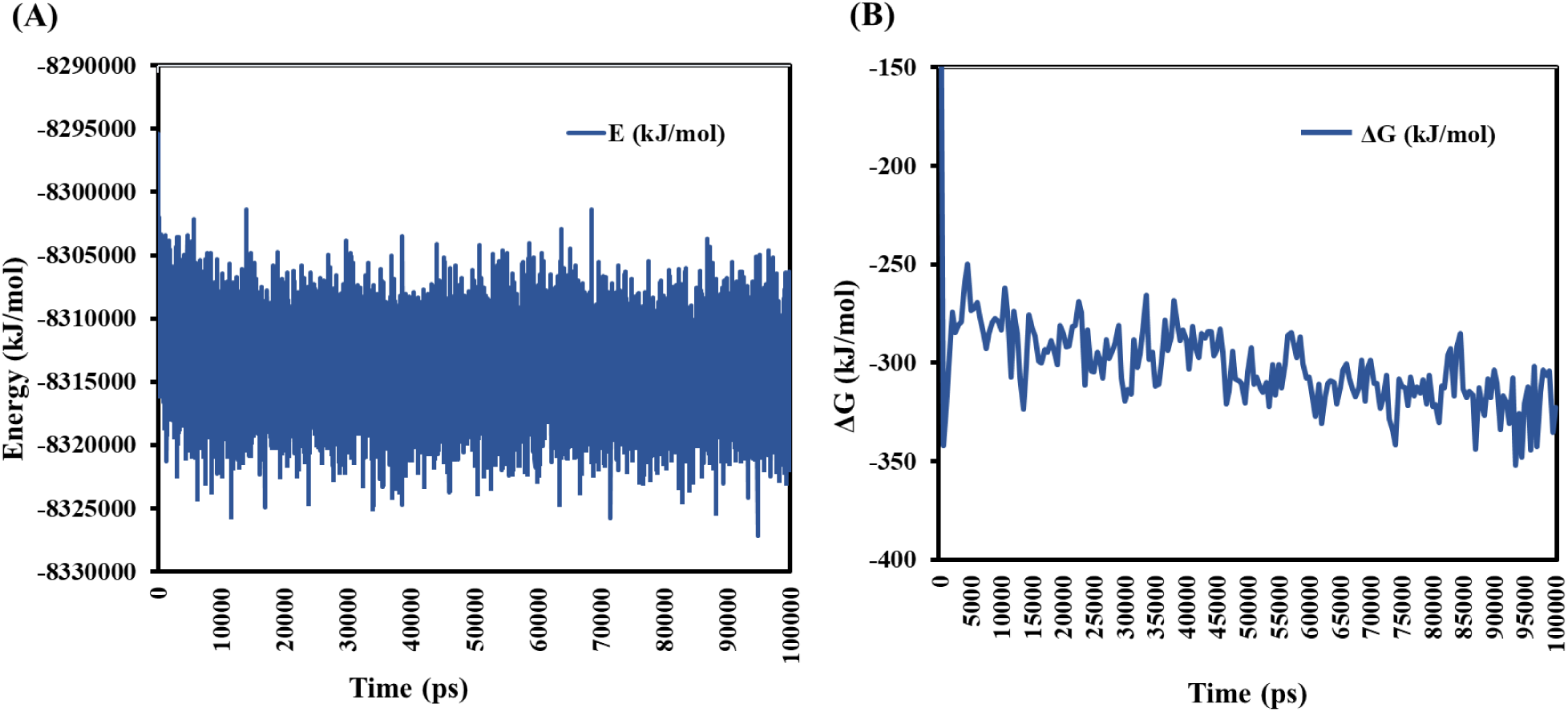
Binding energy analysis of the yTop1–Rpc82 complex during molecular dynamics simulation (A) Time-resolved energy plot of the yTop1–Rpc82 complex over the 100 ns simulation trajectory, showing stabilization of binding energy early in the simulation. (B) MM-PBSA–based binding free energy calculation of the interaction interface, confirming the energetically favorable nature of the complex.

## 4. DISCUSSION

In this study, a full-length structural model of yeast Topoisomerase I (yTop1) was generated using a domain-based modeling approach, which successfully resolved regions, particularly the N-terminal domain (D1) that was previously disordered in AlphaFold and homology-based predictions. We believe that this alternative approach may successfully address the challenges of predicting the three-dimensional structure of such large and highly flexible proteins. Multiple advanced modelling tools were employed to maximize the number of initial models of individual domains in an unbiased manner. Selection of a model for each segment was the outcome of extensive steps of validation using ERRAT, Verify3D, and Ramachandran plot analyses. These measures provided confidence in the reliability of our predicted structure. Modeller program was employed to assemble selected models for individual segments into a full-length structure. Confidence for acceptance of the final model was enhanced due to the application of rigorous structural validation and screening by comprehensive metrics. Proper folding and association of segments in the full-length structure revealed some degree of flexibilities at the N-terminal domain and linker domain regions. However, the other three domains of Topoisomerase I, namely DNA binding domain, Core and C-terminal domain harbouring the catalytic site, were found to have stable modular organization, in agreement with their involvement in the catalytic cycle of the enzyme. Superimposition of the predicted yTop1 model onto the AlphaFold structure revealed minimal deviations, while comparison with the human Top1 crystal structure (PDB ID: 1K4T) showed close alignment in conserved regions, particularly around the active site and DNA-binding interface. These findings support the accuracy of the predicted model and provide a robust foundation for subsequent computational analyses.

Molecular docking studies with Rpc82 protein followed by molecular dynamic simulations aided in evaluating the yTop1- Rpc82 interaction surface under different conditions. Hence, the present study describes an approach for the development of 3D-structural model of full-length Topoisomerase I, which might be a reliable alternative. This model for full length Topoisomerase I would help to further predict its interaction interfaces, model its binding with other essential cellular proteins in order to decipher its role in vital cellular processes in budding yeast. As topoisomerases are known to play critical role in DNA transcription, we further explored molecular interaction between yeast Topoisomerase I (yTop1) with RNA pol III subunit Rpc82 as they were shown to interact previously in high throughput protein-protein interaction studies [25], [26]. The molecular docking results between full-length yTop1 and Rpc82 protein indicated the primary involvement of N-terminal domain (D1) as well as residues belonging to the core-domain (D3) of yTop1. Interestingly, Top1-NTD is often implicated in regulatory interactions. It has been shown to be involved in interaction with RNA pol II, Sir2 [15]. Thus, D1 domain may serve as a flexible docking platform with surface-exposed motifs or conserved patches serving as hotspots for interaction by different partners of Top1 like Rpc82, depending on cellular context. D3 core domain of yTop1 has been previously shown to interact with Tof1, a component of the fork protection complex. yTop1-Tof1 interaction plays a crucial role in replication fork pausing at programmed pause sites like the ribosomal DNA (rDNA) replication fork barrier [98].

Although our intial docking analysis revealed two distinct binding surfaces on Rpc82, the binding interface comprising the WH2–WH3 domains was prioritized for further analysis. This interaction is unlikely to interfere with RNA Pol III function, as yTop1 preferentially binds to Rpc82’s WH2–WH3 domains, which are spatially distinct from the holoenzyme assembly interface. Instead, this association may enhance transcriptional efficiency of RNA polymerase III by facilitating the targeted recruitment of yTop1 to actively transcribing loci. Topoisomerase I plays a critical role in resolving transcription-induced DNA supercoiling, and its proximity to RNA Pol III could enable rapid relaxation of torsional stress, thereby supporting uninterrupted elongation. Moreover, previous studies focusing on RNA polymerase - Topoisomerase I physical interaction till date, demonstrated a positive impact of this interaction in maintaining transcriptional fidelity and genome stability. RNA Pol III is capable of efficiently reinitiating transcription at downstream promoters, a process that is enhanced by the presence of Topoisomerase I. In human systems, Topoisomerase I has been shown to promote multiple rounds of RNA Pol III–mediated transcription from preformed preinitiation complexes [24], suggesting a functional synergy between the two enzymes. Topoisomerase I alleviates transcription-induced supercoiling, thereby facilitating rapid reinitiation and sustained transcriptional output. The physical association described here between yeast RNA Pol III and Topoisomerase I may serve a similar role. By resolving torsional stress at actively transcribing loci, yTop1 could enable uninterrupted rounds of RNA pol III mediated transcription, enhancing throughput and maintaining genomic stability under dynamic cellular conditions.

Comparative analysis of the yTop1–Rpc82 complex with known heterodimers underscores the robustness of the docking methodology and supports the accuracy of the predicted interaction. Strong agreement with benchmark protein complexes, along with consistent binding energy, RMSD, and SASA profiles, confirms the structural and energetic feasibility of this interaction under physiological conditions. Stable RMSD values and moderate RMSF at the interface further highlight the biological relevance of the selected binding surface. MD simulations revealed persistent contacts between specific residues, identifying key interaction hotspots critical for complex stability. Rpc82’s physical association with yTop1 may facilitate its recruitment to actively transcribing RNA Pol III loci, aiding in the resolution of DNA supercoiling and promoting transcriptional fidelity. Rpc82 subunit has been shown to be important component of RNA Pol III holoenzyme with specified role during transcription initiation [95]. Rpc82 via its physical interaction with yTop1 may facilitate recruitment of yTop1 to the sites harbouring active RNA Pol III transcribing genes. Additionally, it might play a critical role in the maintenance of genomic stability by ensuring smooth progression of transcription via its interaction with Topoisomerase I. A stable interaction between these two proteins ensures efficient coordination of these two processes i.e. maintaining genome stability and transcriptional fidelity. MD simulation studies also revealed amino acids residues maintaining persistent interaction between these two proteins and led to identification of key amino acid hot spot residues on the surface, suggesting their critical role in mediating these interactions. This study opens up avenues for further investigation on functional characterization of this PPI and exploring the physiological relevance of RNA pol III- Topoisomerase I connection.

## 5. CONCLUSION

Topoisomerase I in yeast plays a critical role in maintaining DNA topology during essential cellular processes and contributes to genome stability. However, the absence of a well-defined full-length structure has limited structure-based interactome studies and hindered insights into its functional partnerships. This study presents a robust structural framework for full-length yTop1, developed through a domain-based modeling strategy and validated using multiple computational tools, offering a reliable platform for future structural and functional investigations.

The integration of advanced modeling and validation techniques enabled accurate reconstruction of large multidomain proteins such as Topoisomerase I, while molecular dynamics simulations revealed a stable and biologically plausible interaction between yTop1 and the Rpc82 subunit of RNA polymerase III. Identification of key interaction hotspots further strengthens the case for a functional association, suggesting a coordinated role of this physical interaction in transcriptional regulation and genome maintenance. Results of the interaction hotspot analysis lay the groundwork for future experimental validation and targeted mutagenesis studies, opening new avenues to explore the physiological relevance of the Top1–Rpc82 interaction and deepening our understanding of the dynamic interplay between transcription and DNA topology in yeast cells. Importantly, the insights gained here may extend beyond yeast biology. As both these enzymes are conserved across eukaryotes, their interaction might also be conserved. In that scenario, Rpc82–Top1 interface in human cells may represent a novel regulatory axis with potential relevance in cancer cells where transcriptional stress and genomic instability are hallmarks. This study thus lays the groundwork for future investigations into the therapeutic potential of Topoisomerase I-RNA pol III link in human cancers.

## ACKNOWLEDGMENTS

Infrastructural support (DBT-Builder Grant No. BT/PR11357/INF/22/197/2014, BT/INF/22/SP45088/2022 and DST-FIST grant-SR/FST/LSI-560/2013) from Government of India to Department of Life Sciences, Presidency University Kolkata is sincerely acknowledged. S.C. acknowledges CSIR-IICB for infrastructural and financial support. P.N. is the recipient of senior research fellowship from University Grants Commission, Government of India (Ref No. 211610090847). I.M.K. is the recipient of PhD fellowship (DBT/2019/IICB/1213) from Department of Biotechnology, Government of India.

## FUNDING

This research received no external funding

## COMPETING INTERESTS

The authors have no competing interests to declare that are relevant to the content of this article

## AUTHOR CONTRIBUTIONS

S.S.: conceptualization, methodology, supervision, writing-review & editing, S.C.: conceptualization, methodology, supervision. P.N.: investigation, validation, visualization, writing-original draft. I.K.: methodology development, investigation, validation, editing. All authors read and approved the final version of the submitted manuscript.

## DATA AVAILABILITY

All data generated and analyzed during this study, as well as the computational resources utilized, are publicly accessible. The identification of Topoisomerase I interactors was performed through comprehensive searches in PubMed (https://pubmed.ncbi.nlm.nih.gov/), BioGRID (http://thebiogrid.org/), and STRING (https://string-db.org/) databases. Domain architecture information for yeast Topoisomerase I (yTop1) was obtained from the InterPro server (https://www.ebi.ac.uk/interpro/). Structural models of yTop1 domains were rigorously predicted using a variety of algorithmic means like Ab initio, DeepFold and RoseTTAFold through the Zhang Lab server (https://zhanggroup.org/research/#StructurePrediction) and the Robetta server (https://robetta.bakerlab.org/). The crystallographic structure of yeast Topoisomerase I was retrieved from the RCSB Protein Data Bank (PDB ID: 1OIS; https://www.rcsb.org/structure/1OIS). Structural quality of the predicted models were evaluated by ERRAT, Verify 3D, Ramachandran plot parameters utilising the SAVES v6.1 (https://saves.mbi.ucla.edu/) server. Assembly of individual yTop1 domains was performed by the MODELLER software package (https://salilab.org/modeller/). Molecular docking analyses were conducted using three independent servers, ensuring the robustness of our findings: ClusPro (https://cluspro.org/), HDOCK (http://hdock.phys.hust.edu.cn/), and HADDOCK 2.4 (https://rascar.science.uu.nl/haddock2.4/). Interaction parameters including binding energy, interaction surface area etc. were estimated using PDBePISA (https://www.ebi.ac.uk/pdbe/pisa/) and PRODIGY (https://rascar.science.uu.nl/prodigy/) servers. For preparing standard graphs of binding energy and energy/surface area of published homodimers, protein information were collected from RCSB-PDB database (https://www.rcsb.org/structure/), interacting surface area were measured using PDBsum (https://www.ebi.ac.uk/thornton-srv/databases/pdbsum/Generate.html) program and binding energy were evaluated by PRODIGY (https://rascar.science.uu.nl/prodigy/) server. Structural visualization and superimposition of protein structures were achieved by utilizing the Biovia Discovery Studio 2024 free version (https://discover.3ds.com/discovery-studio-visualizer-download), UCSF Chimera X 1.19 (https://www.cgl.ucsf.edu/chimerax/cgi-bin/secure/chimerax-get.py?file=1.9/windows/ChimeraX-1.9.exe) and SuperPose program (https://superpose.wishartlab.com/). Binding free energy calculations were performed using the gmx_MMPBSA package (https://github.com/Valdes-Tresanco-MS/gmx_MMPBSA). Raw data files, including model prediction, docking results, molecular dynamics simulation using installed Gromacs (https://www.gromacs.org/) program and interaction analyses, are available from the corresponding author and archived at Zenodo (https://zenodo.org/records/15622372).

## Notes

https://zenodo.org/records/15622372

### Competing Interest Statement

The authors have declared no competing interest.

