## Supplementary Information for "Computational insights into the interaction between Topoisomerase I and Rpc82 subunit of RNA Polymerase III in *Saccharomyces cereviseae*"

**SUPPORTING INFORMATION**

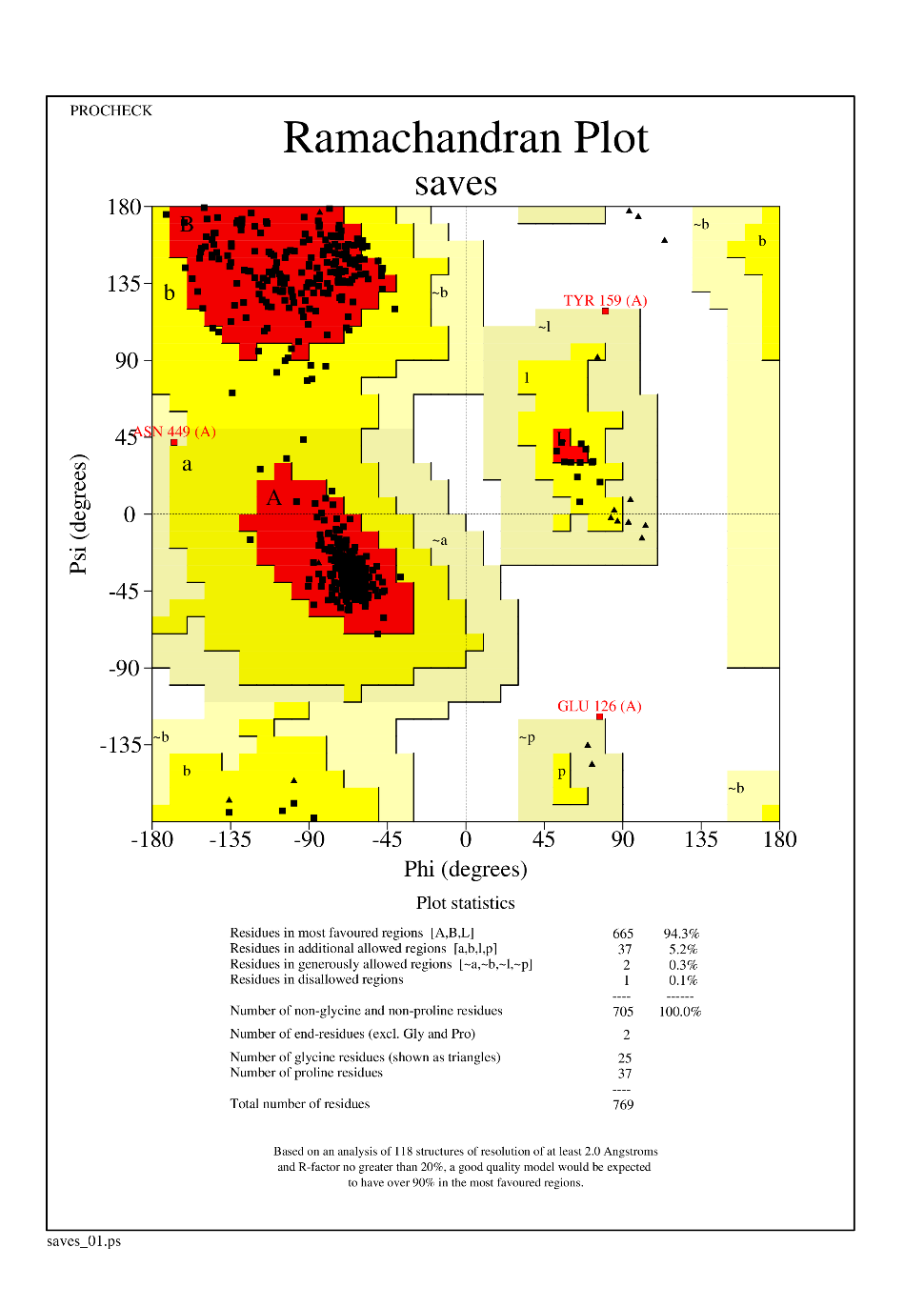

**Supplementary Figure 1: Ramachandran plot for the predicted model of the full-length Topoisomerase-I from budding yeast, indicating the presence of maximum residues at the favoured regions.**

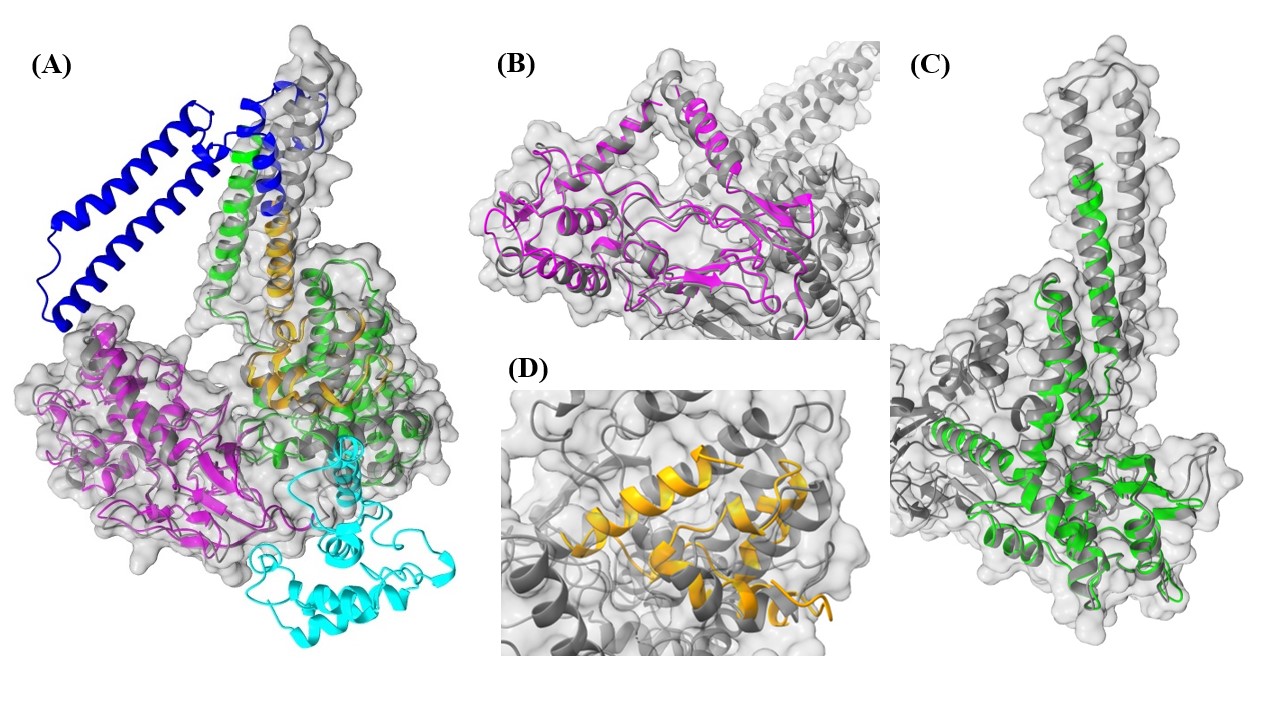

**Supplementary Figure 2: Superimposed image of yeast Topoisomerase I predicted model on human Topoisomerase I X-ray crystallographic structural model (coloured by grey surface with grey ribbon). A- Different domains of full-length predicted yeast Topoisomerase I coloured differently (D1-cyan, D2-magenta, D3-green, D4-blue and D5-orange) and superimposed on human Topoisomerase I (grey). B- Enlarged view of superimposed region of D2 (magenta) of yTop I on human Top IMagnified view of yTopI D5 region (orange) superimposed on hTopI**. C- **superimposed region of D3 (green) of yTop I on human Top I 9grey). D-Magnified view of yTopI D5 region (orange) superimposed on hTopI.**

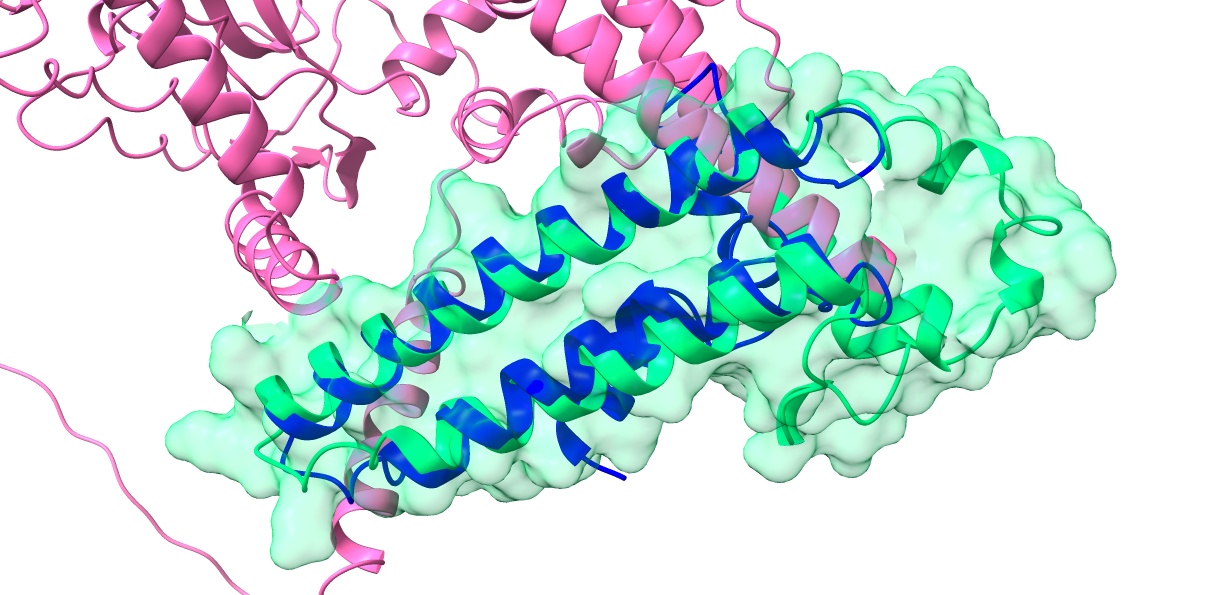

**Supplementary Figure 3: Superimposed image of predicted linker region (D4) of yeast Topoisomerase I (Blue) onto the same region, i.e. spanning from 590-698 amino acid residues (green) from the AlphaFold predicted structural model of full-length yeast Topoisomerase I (pink).**

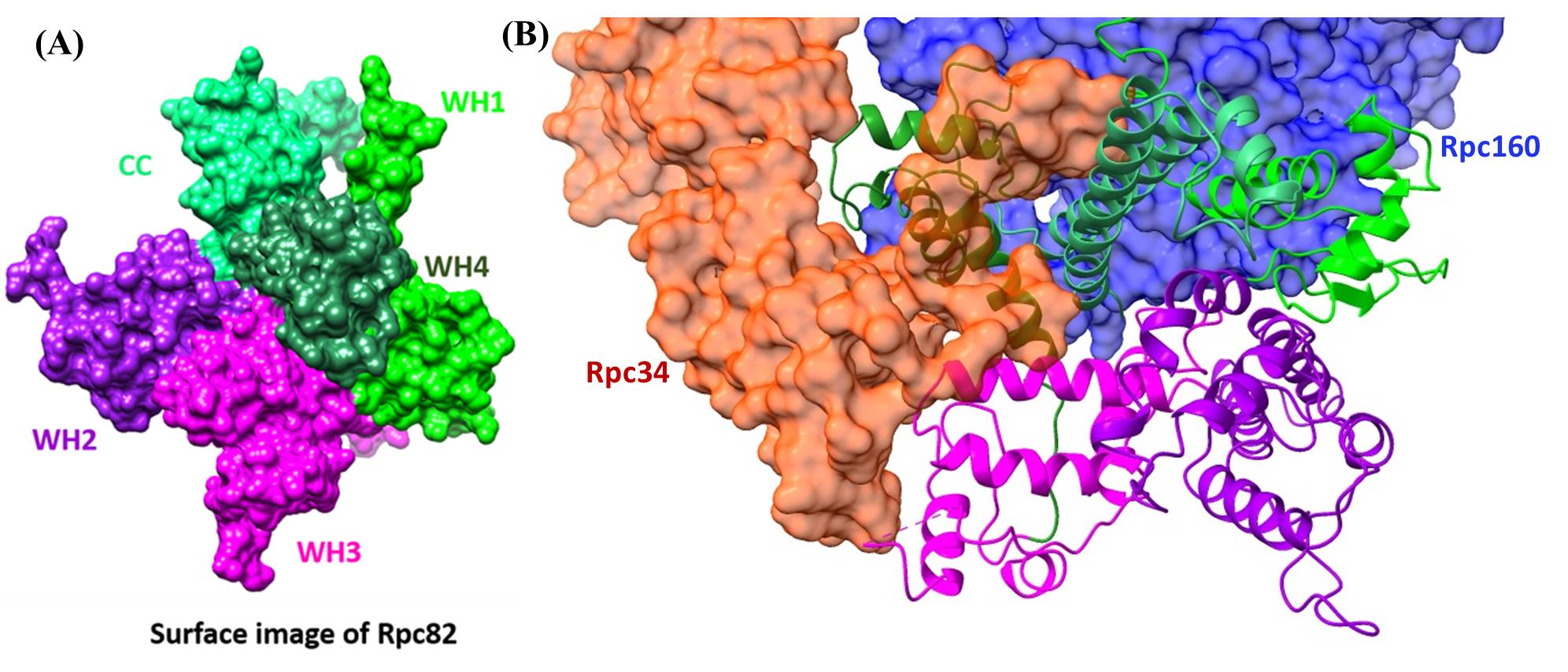

**Supplementary Figure 4: structural assembly analysis of RNA Pol III holoenzyme (PDB ID: 6EU0) revealed that Rpc34 (red) and Rpc 160 (Blue) interacts with the WH1-WH4-CC made surface (green) of Rpc82 protein and the WH2 (Violet) and WH3 (Purple) are unoccupied by any subunits.**

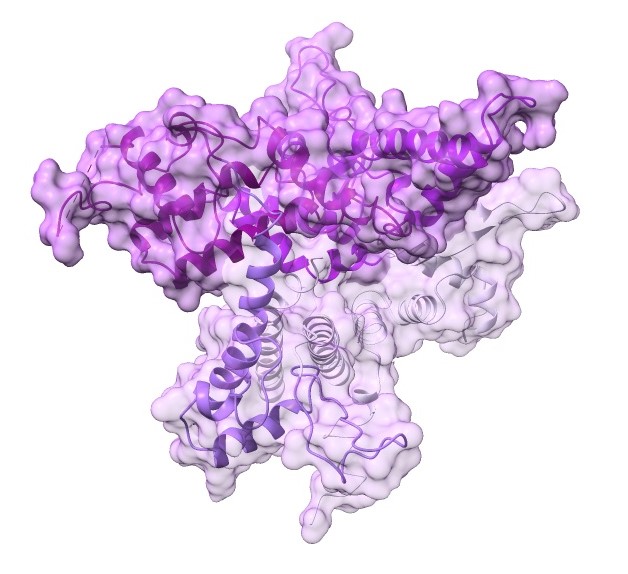

**Supplementary Figure 5: Rpc82 protein with two distinct surfaces, WH1-WH4-CC made surface (lighter shade) and WH2-WH3 made surface indicated by darker shade of violet.**

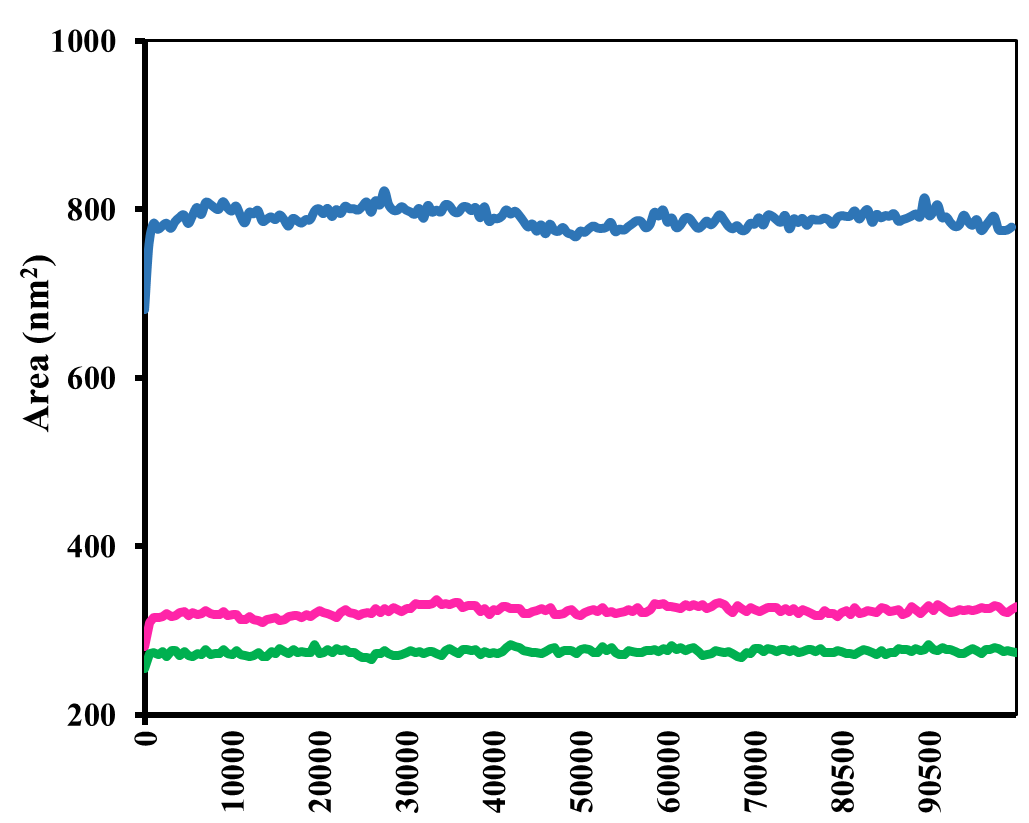

**Supplementary Figure 6: SASA plot of yTop1-Rpc82 complex during the 100 ns simulation (blue) compared with the interaction regions of yTopI (pink) and Rpc82 (green)**

| Parameter | score |
| --- | --- |
| DOPE | -79286 |
| molpdf | 4632.84 |
| ERRAT | 93.43% |
| VerifY3D | 86.35% |
| PROCHECK: |  |
| Ramachandran favoured | 94.30% |
| Allowed region | 5.20% |
| Generously allowed | 0.30% |
| Disallowed region | 0.10% |

**Supplementary Table 1: Different quality assessing parameters and respective score based on which the predicted model for budding yeast Topoisomerase-I was evaluated.**

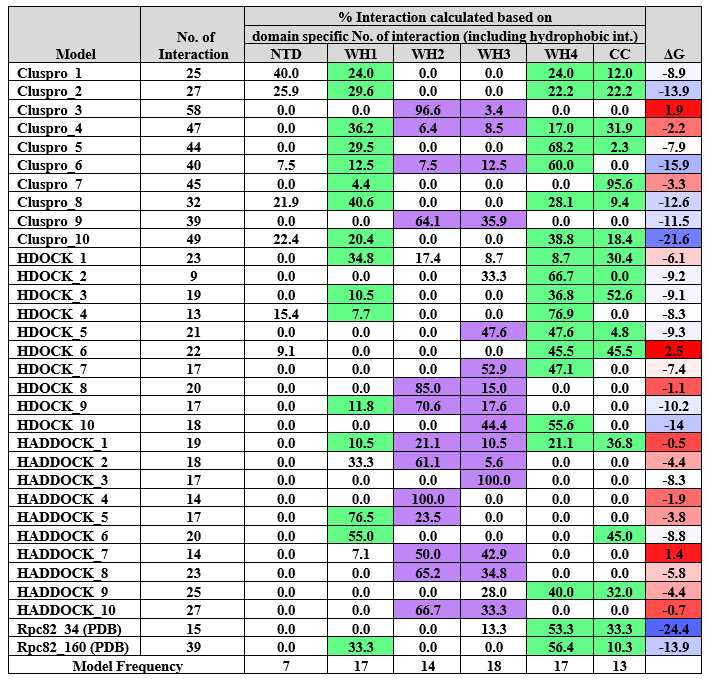

**Supplementary Table 2: Molecular docking interaction analysis for full length yTop I with Rpc82. Ten top scoring models from each server were selected for analysis. Total number of interactions are mentioned in the second column. Interaction of yTop I at different domains of Rpc82 surface was estimated and converted to percentage values to determine the contribution of each domain of Rpc82 towards the total binding interactions observed with yTop I. In few models, yTop I was observed to bind at WH1-WH4-CC surface, which are shaded in green colour whereas models wherein WH2-WH3 surface of Rpc82 formed the binding surface are shaded in violet color. Column with ΔG values have also been shaded with different colours, where blue indicated better (lowest values) binding free energies and red indicated higher values. Interaction between Rpc82-Rpc34 and Rpc82-Rpc160 were also measured (collected from PDB id 6EU0) and mentioned at final two rows**

| yTop1 domain | ΔG value (kcal/mol) | Interaction surface area (Å^2^) | Preferred docking site at Rpc82 |
| --- | --- | --- | --- |
| D1 | -14.2 | 1646.4 | WH2 and WH3 |
| D2 | -13.6 | 1440.5 | Mainly at WH2 and limited WH3 |
| D3 | -15.5 | 1915.5 | WH2, WH3 and few residues of WH1 |
| D4 | -13.2 | 1392.5 | WH2 and WH3 |
| D5 | -12.7 | 1313.5 | WH2 and WH3 |

**Supplementary Table 3: Result analysis of the domain wise targeted docking of yTop I with Rpc82 . All the individual domains of yTop I mostly prefer to dock at the Rpc82 surface formed by WH2 and WH3 domains.**

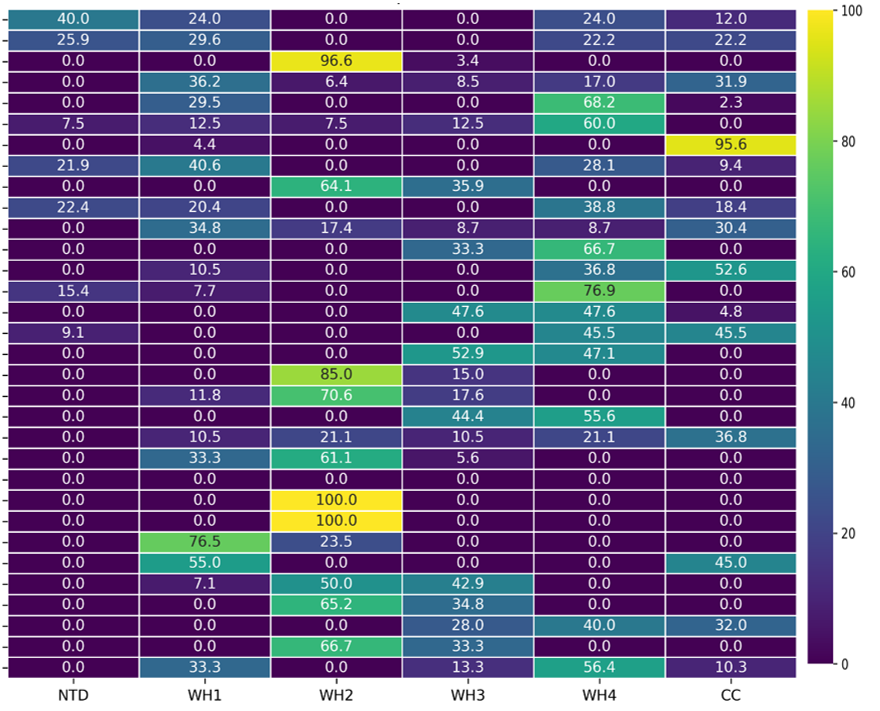

**Supplementary table 4: Heatmap analysis to identify which domains of Rpc82 interact preferentially with different docking models of yTop1, using color gradients to determine the interaction intensity. It provides percent interaction values across different domains of Rpc82 (NTD, WH1-WH4 and CC) for the different docking models involving full-length yTop1. Interaction data for Rpc82’s NTD, WH domains (WH1, WH2, WH3, WH4) and the coiled-coil (CC) domain with full-length yTop1 from docking simulations were gathered and structured in a format that includes domains as columns and docking models as rows, with the cell values representing the percentage of interaction observed in the model. Data from 30 models have been arranged sequentially with 10 best models each from ClusPro, HDOCK and HADDOCK server respectively.**

| **Sl. No.** | **PDB ID** | **Binding energy** | **kcal/mol/Å^2^** |
| --- | --- | --- | --- |
|  |  | **kcal/mol** | **Energy/Surface Area** |
| 1 | 1A22 | -15.7 | -0.012 |
| 2 | 1DJ7 | -10.7 | -0.014 |
| 3 | 1E44 | -10.9 | -0.009 |
| 4 | 1H2V | -12.3 | -0.010 |
| 5 | 1H32 | -11.4 | -0.009 |
| 6 | 1KA9 | -11.3 | -0.008 |
| 7 | 1OF5 | -12.5 | -0.009 |
| 8 | 1OO0 | -10.5 | -0.009 |
| 9 | 1UGH | -15.3 | -0.014 |
| 10 | 1WPX | -13.5 | -0.008 |
| 11 | 2APO | -9.6 | -0.008 |
| 12 | 2B42 | -13 | -0.010 |
| 13 | 2D74 | -11.3 | -0.008 |
| 14 | 2OMZ | -17.3 | -0.013 |
| 15 | 2P1M | -11.6 | -0.007 |
| 16 | 2RAW | -7.4 | -0.005 |
| 17 | 2VSM | -12 | -0.009 |
| 18 | 2XFG | -14.2 | -0.013 |
| 19 | 2Z5B | -12.2 | -0.011 |
| 20 | 2Z64 | -13.6 | -0.012 |
| 21 | 2ZSI | -11.4 | -0.009 |
| 22 | 3AWU | -12.6 | -0.011 |
| 23 | 3CPT | -12 | -0.010 |
| 24 | 3CX8 | -12.9 | -0.009 |
| 25 | 3H7H | -12.4 | -0.008 |
| 26 | 3IEY | -11.6 | -0.008 |
| 27 | 3KXC | -8 | -0.008 |
| 28 | 3MXN | -11.9 | -0.009 |
| 29 | 3N4I | -12 | -0.009 |
| 30 | 3SHG | -11.7 | -0.008 |
| 31 | 3ZYI | -11.1 | -0.011 |
| 32 | 4BL7 | -10.6 | -0.008 |
| 33 | 4CBU | -10.7 | -0.010 |

**Supplementary Table 5:** **Established Heterodimeric proteins of similar size and interaction surface area compared with the selected Top1-Rpc82 complex in terms of binding energy. Data were collected from RCSB-PDB database. Binding energies were calculated using Prodigy server to prepare the standard plot.**

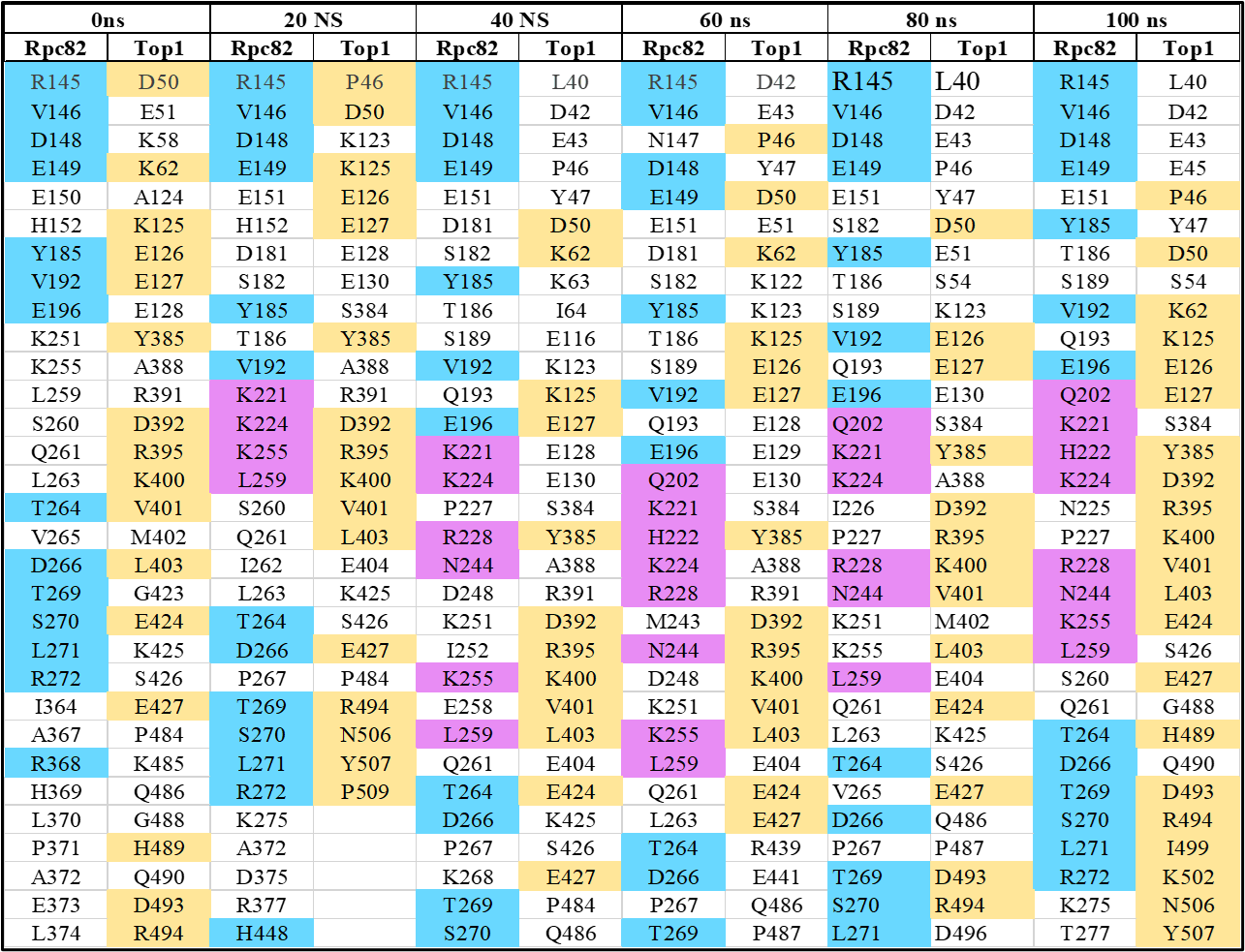

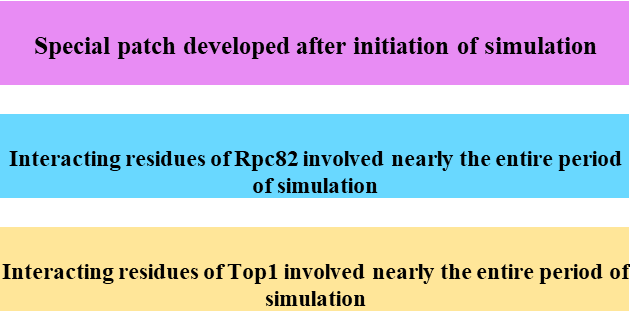

**Supplementary Table 6: Identification of key amino acid residues forming the binding interface between the two proteins (yeast Topoisomerase I and Rpc82 subunit of RNA polymerase III). These residues seem to be involved in forming the binding interface between the two proteins throughout the simulation period. The amino acid residues of yTop I, which seem to maintain contact with the partner protein Rpc82 almost throughout the simulation period have been highlighted in yellow. On the other hand, amino acid residues of Rpc82, which seem to maintain contact with the partner protein yTop I almost throughout the simulation period have been highlighted in blue. Interacting residues that emerge during the course of simulation have been depicted in violet color.**

**All data supporting the findings of this study are available in a Zenodo repository and accessible through Zenodo DOI: . (**[**https://zenodo.org/records/15622372**](https://zenodo.org/records/15622372)**).**
